# Sequencing a botanical monument: a chromosome-level assembly of the 400-year-old Goethe’s Palm (*Chamaerops humilis* L.) at the Botanical Garden of the University of Padua (Italy)

**DOI:** 10.1101/2025.03.17.643598

**Authors:** Núria Beltran-Sanz, Stefan Prost, Veronica Malavasi, Silvia Moschin, Carola Greve, Tilman Schell, Tomas Morosinotto, Francesco Dal Grande

## Abstract

The rapid decline in global biodiversity highlights the urgent need for conservation efforts, with botanical gardens playing a crucial role in *ex situ* plant preservation. Monumental plants, such as the 400-year-old Goethe’s Palm (*Chamaerops humilis* L.) at the UNESCO Botanical Garden of the University of Padua (Italy), serve as vital flagship species with significant ecological and cultural value. In this study, we present the first high-quality, chromosome-level genome assembly of *C. humilis*, using PacBio HiFi and Arima Hi-C sequencing technologies. This genome is the most contiguous and complete within the Arecaceae family to date, with an exceptionally high repeat content of 88%, of which 63% is attributed to Long Terminal Repeat (LTR) elements. Comparative analysis of the transposable element (TE) landscape in palms suggests that the LTR expansion in *C. humilis* is likely the result of recent TE bursts. Furthermore, we provide the first comprehensive annotation of microRNAs (miRNAs) in the Arecaceae family, identifying, for the first time in palms, the miRNA family miR827, which plays a key role in nutrient regulation. Microsatellite analysis suggests that it probably belongs to the Western genetic lineage of *C. humilis*. These findings represent a significant advancement in the conservation genomics of this species, laying the groundwork for enhanced preservation strategies, with botanical gardens as pivotal actors in the fight to safeguard global biodiversity.

## Introduction

Biodiversity is globally facing a rapid decline, necessitating urgent conservation measures to preserve plant diversity. Botanical gardens worldwide are pivotal in this effort, conserving an estimated 100,000 plant species *ex situ* (Mounce et al., 2017). Beyond their roles in recreation, education and science communication, these gardens function as vital centers for conservation and research, implementing integrated programs to prevent plant extinctions (Ren & Antonelli, 2023). The *ex situ* conservation approach of botanical gardens involves maintaining and cultivating plants outside their natural habitats, which acts as a safeguard against the total loss of species in the wild. This includes a diverse array of activities such as seed banking, tissue culture, and creating living collections that serve as genetic reservoirs. These efforts are crucial, especially in light of rapid habitat destruction and climate change impacts that threaten many plant species (Heywood, 2017).

Monumental trees—large, ancient, or historically significant plants that are notable for their size, age, rarity, or cultural importance (Cannizzaro & Corinto, 2014), serve as effective flagship species for conservation initiatives led by botanical gardens. These trees are not merely ecological relics; they are also invaluable cultural and educational assets (Vannuccini et al., 2006; Cannizzaro & Corinto, 2014). Their iconic appeal makes them powerful symbols for conservation campaigns, effectively garnering support from diverse stakeholders. Their preservation not only protects genetic diversity but also maintains cultural heritage, offering educational opportunities that bridge the gap between science and society (Zapponi et al., 2017; Schicchi et al., 2021). In addition, monumental trees that are known to have lived for hundreds of years could also represent interesting models to study long-term adaptation responses.

In the field of conservation genomics, these ancient plants can play a crucial role in the new era of reference genomes (Formenti et al., 2022) by bridging the gap between modern genomic techniques and traditional conservation efforts. Advancements in sequencing technologies have dramatically reduced costs and time, allowing for comprehensive genetic mapping of many species (Formenti et al. 2022; Li et al. 2022). The detailed genetic maps obtained from sequencing monumental trees can serve as reference genomes, providing a baseline for future genetic studies and conservation efforts and underscoring the importance of genetic research in conservation efforts (Theissinger et al., 2023). It will also allow researchers to uncover how they have adapted to survive through centuries of environmental changes. For example, the genome sequencing of the Napoleon Oak (*Quercus robur* L.) in Switzerland revealed remarkable genetic stability, with low levels of somatic mutations even in an ancient tree (Schmid-Siegert et al., 2017). This suggests robust mechanisms for maintaining genetic integrity, essential for the oak’s long lifespan. Similarly, the Olive tree (*Olea europaea* L.) of Vouves in Greece, estimated to be between 3,000 and 5,000 years old, provided critical genetic insights through genome sequencing (Bombarely et al., 2021). Researchers identified different genetic origins in the tree’s rootstock and scion, indicating ancient grafting practices.

In this study, we present the first high-quality genome assembly of *Chamaerops humilis* L. (Arecaceae) by sequencing Goethe’s Palm, the oldest individual specimen at the UNESCO Botanical Garden of the University of Padua (Italy) and the oldest cultivated palm in the world (Hodge, 1982). This specimen holds significant value for several reasons.

First, from a scientific and biogeographical perspective, Goethe’s Palm is crucial for understanding the adaptive traits that allow this species to thrive in Mediterranean climates, which differ significantly from the tropical regions typically associated with palm species. This makes *C. humilis* stand out as the palm species occurring at the northernmost latitudes (Garcia-Castano et al., 2014). Previous research using ten microsatellites identified two distinct genetic lineages within the species: one predominantly found in the western Mediterranean, including populations in Spain and Portugal, and the other in the eastern Mediterranean, encompassing populations in southern Italy and Tunisia, which are characterized by lower genetic variability (Giovino et al., 2023).

Second, historically, Goethe’s Palm is a cultural and scientific landmark. During his visit to Padua in 1786, the polymath and writer Johann Wolfgang von Goethe was inspired by this palm to develop his foundational botanical concepts featured in his scientific book, *The Metamorphosis of Plants* (von Goethe, 1790). He observed that all plant organs, such as leaves, petals, and stems, were modifications of a basic, idealized form (the “Urpflanze” or “archetypal plant”), and he found the leaves of the palm at Padua’s Botanical Garden particularly fascinating – so much so that he kept samples of it for the rest of his life (Miller, 2009). Sequencing the genome of Goethe’s famous palm may shed light on its provenance as it remains unclear where the specimen originated from when it was first cultivated in 1585, and this insight may deepen the historical significance of this treasure.

Third, from a conservation perspective, *C. humilis* is under threat despite being listed as ‘Least Concern’ by the IUCN Red List (IUCN 2024). The species faces pressures from habitat loss, urbanization, and agricultural expansion, leading to declining populations in the wild (Martínez-Vega & Rodríguez-Rodríguez, 2022; Nekhla et al., 2022; Giovino et al., 2023). As a Pleistocene relict species and a thermo-Mediterranean bioindicator (Herrera, 1989), its conservation is critical for preserving the genetic diversity and ecological history of Mediterranean habitats. By positioning *C. humilis* as a potential flagship species for the conservation of Mediterranean Pleistocene relics, the genetic insights gained from sequencing Goethe’s Palm can serve as a fundamental step towards understanding the species’ adaptive traits and resilience. These findings may be transferable to other species with similar ecologies and biogeographic histories. Specifically, our objectives were three-fold: i) to obtain a high-quality reference genome of *C. humilis* using PacBio HiFi and Arima Hi-C technologies, ii) to compare the genome of *C. humilis* with other available palm genomes to identify and highlight potential genetic peculiarities that could contribute to its unique adaptive traits, and iii) to compare its genetic makeup with the microsatellite variation presented in Giovino et al. (2023) to elucidate the provenance of Goethe’s Palm, gaining insights into its historical origins.

## Materials and Methods

### Genome sequencing

One fresh leaf of Goethe’s Palm (Fig. 1) was collected at the Botanical Garden of the University of Padua (Italy) in September 2022.

**Fig. 1:**
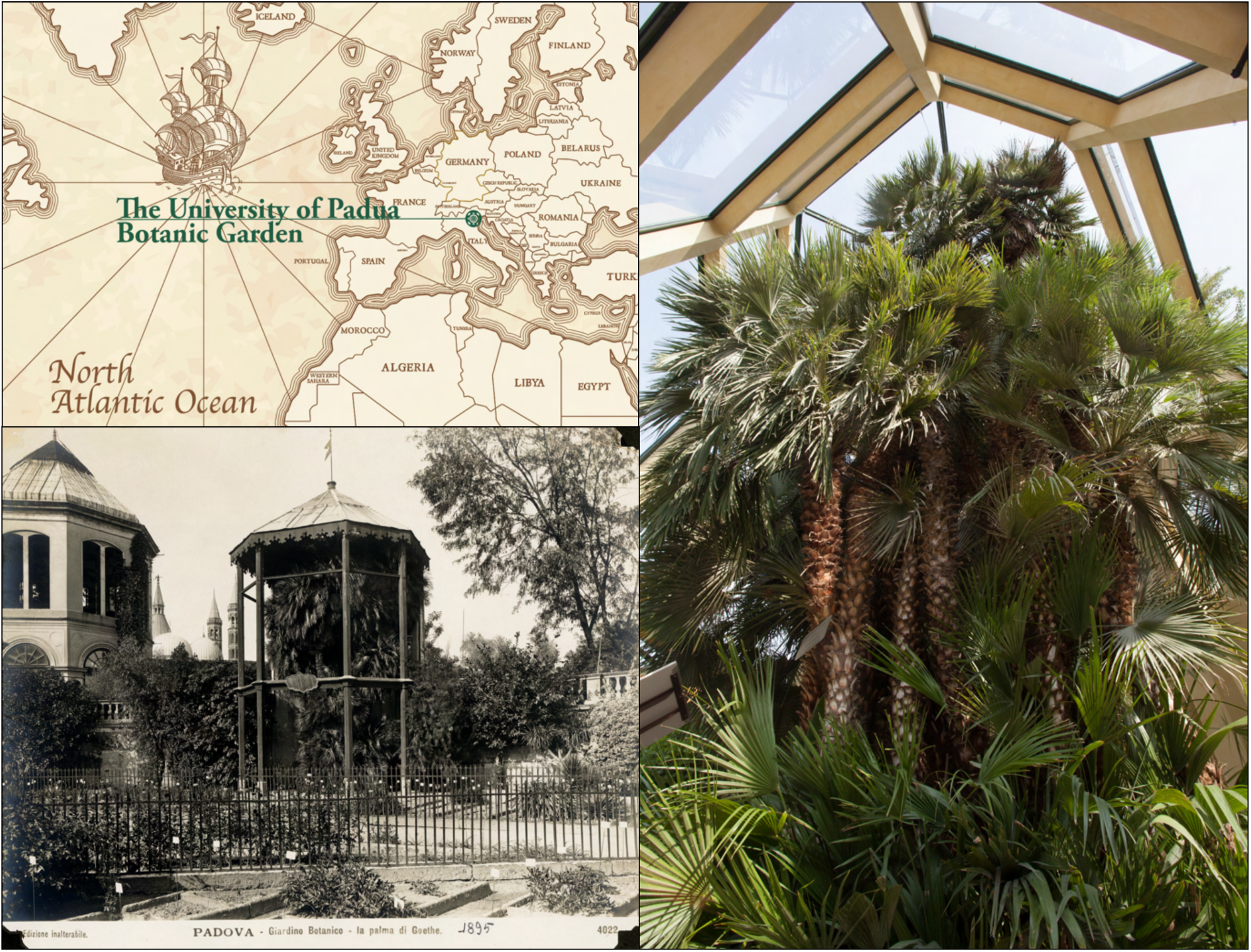
Goethe’s Palm (*Chamaerops humilis*) at the Botanical Garden of Padua (Italy). Top left: location of the Botanical Garden of Padua; bottom left: Goethe’s Palm in 1895; right: the plant as it appears today.

After 5 days in the dark, genomic DNA was extracted from approximately 30 mg of leaf tissue according to the CTAB-based protocol from Murray and Thompson (1980). DNA concentration and DNA fragment length were assessed using the Qubit dsDNA BR Assay kit on the Qubit Fluorometer (Thermo Fisher Scientific) and the Genomic DNA Screen Tape on the Agilent 2200 TapeStation system (Agilent Technologies), respectively. One SMRTbell library was constructed following the instructions of the SMRTbell Express Prep kit v2.0. The same library was loaded twice on the PacBio by performing two SMRT cell sequencing runs on the Sequel System IIe in CCS mode. We then employed a Hi-C approach to obtain high-resolution data on the spatial organization of the genome, which is essential for precise genome scaffolding. The Hi-C library was prepared from approximately 250 mg of leaf tissue (from the same individual) using the Arima High Coverage Hi-C Kit v01 (Arima Genomics, Carlsbad, CA, USA) according to the Animal Tissue User Guide. This kit captures the organizational structure of chromatin in three dimensions by first fixing the chromatin structure using formaldehyde, then digesting the crosslinked chromatin using a restriction enzyme cocktail optimized for coverage uniformity across a wide range of genomic sequence compositions, and finally ligating the ends of the molecules in close proximity using a biotin-labeled bridge. Library preparation of the proximally-ligated DNA was then performed according to the Swift Biosciences Accel-NGS 2S Plus DNA Library Kit protocol. Fragment size distribution and concentration of the Arima High Coverage Hi-C library were assessed using the TapeStation 2200 (Agilent Technologies) and the Qubit Fluorometer and Qubit dsDNA HS Reagents Assay Kit (Thermo Fisher Scientific, Waltham, MA), respectively. The library was sequenced on an Illumina NovaSeq 6000 platform at Novogene (Cambridge, UK) using a 150 bp paired-end sequencing strategy with an insert size of 350 bp.

### RNA sequencing

Four different tissues (young and old leaves, marrow, fruits, and roots) of *C. humilis* were collected for RNA extraction. The microRNA and mRNA were extracted using CTAB extraction buffer (Chang et al., 1993) and subsequently quantified with an Implen NanoPhotometer (Implen GmbH, Munich, Germany). Only the samples with 260/280 and 260/230 absorbance ratios above 1.8 value were kept. For each sample, 10 µg of total RNA was treated with DNase I (New England Biolabs; Ipswich, Massachusetts, USA) to remove contaminant DNA and purified with an RNA Clean & Concentration-5 kit (Zymo Research, California, USA).

The RNA extracts were then pooled, and the library preparation for both microRNAs and mRNA, as well as the sequencing, were performed at Novogene (UK). Sequencing was performed on the NovaSeq 6000 platform (Illumina, Inc., San Diego, CA, USA), generating 150 bp paired-end reads with an insert size of 350 bp.

### Nuclear genome assembly and scaffolding

High-quality circular consensus sequences based on the PacBio subreads were generated using a workflow containing DeepConsensus version 0.2.0 (Baid et al., 2023). Briefly, all CCS reads were obtained from subreads using PacBio’s ccs tool version 6.4.0 (https://github.com/PacificBiosciences/ccs). Both the CCS reads and subreads were then aligned with actc version 0.3.1 (https://github.com/PacificBiosciences/actc). DeepConsensus was run using the CCS reads and alignments. The long high-fidelity (HiFi) fragments generated were first trimmed from potential adapters with HiFiAdapterFilt version 2.0.1 (Sim et al., 2022) and subsequently assembled at the contig level with Hifiasm version 0.18.7 (Cheng et al., 2022).

We scaffolded the primary and haplotype contig-level assemblies with Hi-C data to generate chromosome-level assemblies. This was carried out using the Arima Hi-C mapping pipeline (Arima Genomics 2023). In brief, the Hi-C reads were first aligned to the contigs using BWA-MEM version 0.7.17 (Li, 2013). The 3′ ends of the reads identified as chimeric were then removed with Arima in-house perl script (part of the Arima Hi-C mapping pipeline). Next, duplicated reads were removed with Picard version 3.0.0 (Picard Toolkit 2019). Finally, the assembly was scaffolded to chromosome level using YaHS version 1.1 (Zhou, 2023) and visualized for quality assessment with Juicebox version 1.11.08 (Robinson et al. 2018). To fill gaps, TGS-GapCloser version 1.0.1 (Xu et al., 2020) was run with Racon version 1.4.3 (Vaser et al., 2017) as a method for error correction. The quality of the primary and the two haplotype assemblies was evaluated with Quast version 5.0.2 (Gurevich et al., 2013), Assemblathon2 (Bradnam et al., 2013) and Qualimap version 2.2.1 (García-Alcalde et al., 2012). To assess their completeness, we ran BUSCO version 5.2.2 (Manni et al., 2021) using the Viridiplantae_odb10 dataset.

To enhance the quality of the genome assemblies, the Arima Hi-C mapping pipeline was run once more, using the output from the initial run as input. The subsequent steps involved gap closing, quality assessment and gene completeness of the genome assemblies, as described above. We then used Blobtools version 1.1.1 (Laetsch and Blaxter, 2017) to identify potential contamination. To do so, the high-quality HiFi reads were mapped against each assembly using Minimap2 version 2.26 (Li, 2021), and the scaffolds of the assembly were blasted against the NCBI Nucleotide database with Blastn version 2.14.0 (McGinnis and Madden, 2004). Finally, the genome assembly statistics were summarised in a snail plot, which was obtained using Blobtoolkit version 4.2.1 (Challis et al., 2020).

To evaluate the synteny between the primary assembly and each haplotype, as well as between haplotypes, we ran JupiterPlot v.3.8.1 (Chu, 2018).

Furthermore, we identified telomeres and centromeres of the primary assembly with quarTeT v.1.2.5 (Lin et al., 2023) and CentrIER v.2.0 (Xu et al., 2024), respectively. The default parameters were used for all functions except for telomere identification, where the -m 10 parameter was employed. The results were then visualised in an ideogram plot using the ‘ggplot2’ R package (Wickham, 2016).

### Organelle genome assembly and annotation

The mitochondrial genome was assembled with MitoHiFi version 3.2 (Uliano-Silva et al., 2023) using *Phoenix dactylifera L.* (family Arecaceae; NCBI: NC_016740) as the most closely related reference genome (see Fang et al. 2012 for more details). All contigs belonging to the mitochondrion were identified from the primary chromosome-level assembly using Blastn version 2.14.0 (McGinnis and Madden, 2004) and removed with Seqtk version 1.3 (Li, 2012).

The chloroplast genome was assembled with ptGAUL version 1.0.5 (Zhou et al., 2022) using *C. humilis* (family Arecaceae; NCBI: ON248747) as the most closely related reference genome (see Yao et al., 2023 for more details). In contrast to the mitochondrial genome assembly, the chloroplast assembly was based on HiFi reads (not the genome assembly) and annotated using the Plastid Genome Annotator script ‘PGA.pl’ (Qu et al., 2019). To identify chloroplast scaffolds in the palm genome assembly, we used Blastn version 2.14.0 (McGinnis and Madden, 2004) with the chloroplast assembly. The scaffolds found were then removed from the genome with Seqtk version 1.3 (Li, 2012).

The best reference genome for both the mitochondria and the chloroplast were identified with the script ‘findMitoReference.py’ (Uliano-Silva et al., 2023). In both cases, the organelle genome assembly quality was estimated with Quast version 5.0.2 (Gurevich et al., 2013), and visualized with OGDRAW version 1.3.1 (Greiner et al., 2019). To create the plot, GeSeq version 2.03 (Tillich et al., 2017) was used, keeping only the best annotations, and setting the BLAT annotation engine (BLATX and BLATN) for mitochondrial annotation and the BLAT (BLATX and BLATN), Chlöe, HMMER and tRNAscan-SE_v2 for chloroplast annotation. In both cases, 95% sequence identity for the proteins, rRNA and tRNA was used. As a reference genome, we used all NCBI RefSeq organelle genomes from the Arecaceae family. After that, we used the GenBank file created by GeSeq as an input of OGDRAW version 1.3.1 (Greiner et al., 2019) to obtain the plot in pdf format.

### Transcriptome assembly and quality assessment

Raw reads from the RNA sequencing data (microRNA and mRNA) were first trimmed with Fastp version 0.23.4 (Chen, 2023) and subsequently aligned to the primary chromosome-level assembly with HISAT2 version 2.2.1 (Kim et al., 2019). Each type of RNA was assembled with StringTie version 2.1.2 (Shumate et al., 2022). The completeness of the transcripts obtained was then evaluated with BUSCO version 5.2.2 (Manni et al., 2021) using the ‘Viridiplantae_odb10’ dataset.

Additionally, the microRNAs were aligned and annotated using ShortStack version 4.0.3 (Axtell 2013) with microRNAanno as a reference dataset (Chen et al., 2021). The results were then plotted in an ideogram generated with the ‘ggplot2’ R package (Wickham, 2016).

### Genome annotation of the primary assembly

For increased accuracy during gene prediction, repeat regions in the primary assembly were first masked using both the *de novo* repeat Viridiplantae library in RepeatModeler version 2.0.5 (Flynn et al., 2020) and the Liliopsida repeat library from RepBase (Jurka et al., 2005) using RepeatMasker version 4.1.6 (Tarailo-Graovac and Chen, 2009). The Liliopsida library was selected because *C. humilis* belongs to this taxonomic group, making it the closest available clade. Given the significant impact of masking on gene prediction, two different masking procedures were applied using RepeatMasker version 4.1.6 (Tarailo-Graovac and Chen, 2009): (i) simple repeats were soft-masked while interspersed repeats were hard-masked, and (ii) all repeats were soft-masked.

The gene annotation was performed with both BRAKER3 (Gabriel et al. 2024) and GeMoMa version 1.9 (Keilwagen et al., 2018) using (i) the mapping file obtained after mapping the mRNA to the primary assembly, (ii) the reference genomes of *Elaeis guineensis* Jacq. (GCA_000442705.1, African oil palm, Singh et al., 2013) and *Phoenix dactylifera* L. (GCA_009389715.1, date palm, Hazzouri, et al. 2019), and (iii) the primary assembly masked. To run BRAKER3, we used the soft-masked genome, whereas for GeMoMA, we used the hard-masked genome, as required by each tool. The output of both tools was then combined with previously generated transcriptomic data to obtain a consensus gene annotation using EVidenceModeler version 2.1.0 (Haas et al., 2008).

We analyzed the completeness of the predicted proteins with BUSCO version 5.2.2 (Manni et al., 2021) utilizing the ‘Viridiplantae_odb10’ data set. The functional annotation of the predicted proteins was conducted by a BLASTP version 2.14.0 (McGinnis and Madden, 2004) search with an E-value cutoff of 10^−6^ against the Swiss-Prot database. Finally, InterProScan version 5.64-96.0 (Jones et al., 2014) was run to annotate gene ontology (GO) terms, domains, and motifs.

To search for homology between the proteins of the primary assembly, the sequence similarity between all protein-coding genes was compared using DIAMOND version 0.9.30 (Buchfink et al., 2021). For a protein sequence, the best five hits that met an E-value threshold of 10^-5^ were reported. The file generated was employed in conjunction with the annotation file to execute MCScanX version 1.0 (Wang et al., 2012) to generate pairwise synteny blocks of proteins. To identify the number and type of duplicated proteins, we ran the function ‘Duplicate_gene_classifier’ from MCScanX version 1.0 (Wang et al., 2012). This script classifies the origins of the duplicate genes in the genome assembly into whole genome / segmental (matching genes in synteny blocks), tandem (consecutive repeat), proximal (located in nearby chromosomal regions but not adjacent) or dispersed (any mode other than segmental, tandem and proximal) duplications. Additionally, the script ‘group_collinear_genes.pl’ was applied to cluster the proteins by connecting collinear proteins until no protein in each group had any collinear proteins outside the group. To visualize the collinear proteins along the chromosomes, the SynVisio software (Bandi and Gutwin, 2020) was used for graphical representation.

### Comparison to other published palm genomes

The genomes of 15 palm specimens belonging to eight different species were downloaded from NCBI, and statistics for each of them (assembly length, N50, and GC content) were calculated with Quast version 5.0.2 (Gurevich et al., 2013, see Table S1). BUSCO version 5.2.2 (Manni et al., 2021) with the ‘liliopsida_odb10’ dataset was used to quantify complete and single-copy BUSCO genes of these palm genomes. Only genomes with more than 50% completeness were selected for the BUSCO-based phylogenomic analysis. The phylogenomic position of *C. humilis* with respect to the other related palm species was determined with ‘BUSCO_phylogenomics.py’ (https://github.com/jamiemcg/BUSCO_phylogenomics). In short, this pipeline aligned the BUSCO sequences with MUSCLE version 5.1 (Edgar, 2022), trimmed them with TrimAl version 1.4.1 (Capella-Gutiérrez et al., 2009), and constructed a maximum-likelihood consensus tree with IQ-TREE version 2.2.6 (Nguyen et al., 2015). The genomes of *Aegilops tauschii* subsp. *strangulata* (GCF002575655.2) and *Zea mays* (GCF902167145.1) were used as outgroups.

To assess LTR expansion in *C. humilis* as a result of recent TE bursts, we calculated the percentage of LTRs for each palm species. For species with multiple genome assemblies, the most complete one was selected based on BUSCO scores (Table S1). The proportion of LTR elements was determined by usinging RepeatMasker version 4.1.6 (Tarailo-Graovac and Chen, 2009) as described above. This method involved the integration of the *de novo* repeat Viridiplantae library from RepeatModeler version 2.0.5 (Flynn et al., 2020) and the Liliopsida repeat library from RepBase. The proportion of LTR elements from the Repatmasker-generated table was used to build a linear regression model to examine its linear relationship with the assembly length. Repeat divergence landscapes were then created following the methodology of Rodriguez and Arkhipova (2023), using TwoBit version 2.0.9 (weng-lab.github.io/TwoBit/), and two Perl scripts (calcDivergenceFromAlign.pl and createRepeatLandscape.pl) from RepeatMasker version 4.1.6 (Tarailo-Graovac and Chen, 2009). The results were presented in the form of a histogram created using the ‘ggplot2’ R package (Wickham, 2016).

We downloaded all protein data available from NCBI for palm species, including *Elaeis guineensis, Phoenix dactylifera*, and *Cocos nucifera.* Furthermore, we also downloaded the KW-346 reference dataset from UniProt, which includes proteins involved in the response to stress. We then functionally annotated the predicted palm proteins using a BLASTP version 2.14.0 search (McGinnis and Madden, 2004) with an E-value cutoff of 10^-6^ against the reference set of stress-related proteins. The resulting data were presented as a barplot, constructed with the ‘ggplot2’ R package (Wickham, 2016).

Finally, the mitochondrial and chloroplast genome assemblies were compared with others available in NCBI. Due to the absence of *C. humilis* mitochondrial genomes in NCBI, we compared our results with those of other species in the Arecaceae family (PP035767, NC_031696, JN375330, MH176158, MG257490, and BK059358). For the chloroplast, we compared our assembly with three others for *C. humilis* (ON248747, NC_029967, and KT312935). For each assembly, the GC content was calculated with Quast version 5.0.2 (Gurevich et al., 2013), and the number of genes was calculated with GeSeq version 2.03 (Tillich et al., 2017), as mentioned above.

### Geographical origin of Goethe’s Palm

A recent study by Giovino et al. (2023) examined the genetic structure of *C. humilis* across its native range using a set of specifically designed genetic markers. The ten highly polymorphic microsatellites identified by Giovino et al. (2023) were used in this study to determine the origin of Goethe’s Palm. To perform this analysis, the microsatellites were extracted from haplotype 1 and haplotype 2 chromosome-level assemblies of *C. humilis* using the tools SeqKit version 0.10.2 (Shen et al., 2016) and BLASTn version 2.14.0 (McGinnis and Madden, 2004). The data were subsequently combined with the Giovino et al. (2023) dataset, which comprised approximately 300 individuals distributed across the native geographic range of *C. humilis*. We constructed a genind object using the ‘adegenet’ R package (Jombart and Ahmed, 2011) and removed duplicated genotypes with the ‘poppr’ R package (Kamvar et al., 2014). The dataset obtained was then used in a Discriminant Analysis of Principal Components (DAPC). The results were visualized using the function ‘compoplot’ from ‘adegenet’ R package (Jombart and Ahmed, 2011), and the ‘ggplot2’ R package (Wickham, 2016).

## Results

### Nuclear genome sequencing and assembly

The genome assembly of *C. humilis* was generated by assembling 69 Gb of PacBio HiFi data, with a read N50 of 14,450 bp and a mean length of 12,812 bp. Hi-C sequencing generated a total of 97.4 Gb of data.

After two iterations of the Arima Hi-C mapping pipeline, the primary and the two haplotypes decreased their total number of scaffolds, increased the scaffold N90, and converged into 18 main scaffolds (Table S2, Table S3, and Table S4). These 18 scaffolds displayed on the contact maps (Fig. S1) are consistent with the findings of Röser (1993), who identified 18 chromosomes pairs in *C. humilis*. The GC content remained the same throughout all the iterations, being 43.8% in all the assemblies (Table S2, Table S3, and Table S4).

All telomeres were identified in the primary assembly, with the exception of chromosome 7, which lacked one. In a similar manner, the presence of all putative centromeres was detected, except for chromosome 10 (Fig. 2A). The primary assembly had a total length of 4.41 Gbp, with 1,986 scaffolds, scaffold N50 of 195 Mbp, and BUSCO completeness of 99.3% (Table 1, Fig. 2B, and Table S2).

**Table 1:** Summary of the main statistics of the primary assembly of *C. humilis*.

| Scaffold-level statistics |  |
| --- | --- |
| Number of scaffolds | 1,986 scaffolds |
| Number of chromosomes | 18 main scaffolds |
| Largest scaffold (bp) | 352,408,672 |
| Total length of the scaffolds (bp) | 4,406,632,840 |
| GC content | 43.8 % |
| N50 | 195,022,367 |
| L50 | 9 |
| BUSCO completeness<br>(viridiplantae_odb, n = 425) | C: 99.3% [S: 88.0% and D: 11.3%],<br>F: 0.5%, M: 0.2% |
| Repetition annotation |  |
| Retroelements | 66.06 % |
| LINEs | 2.85 % |
| LTR elements | 62.8 % |
| Ty1/Copia | 37.7 % |
| Gypsy/DIRS1 | 24.85 % |
| DNA transposons | 3.21 % |
| Others (rolling-circles, small RNA,<br>satellites, simple repeats, and low<br>complexity) | 3.77 % |

**Fig. 2:**
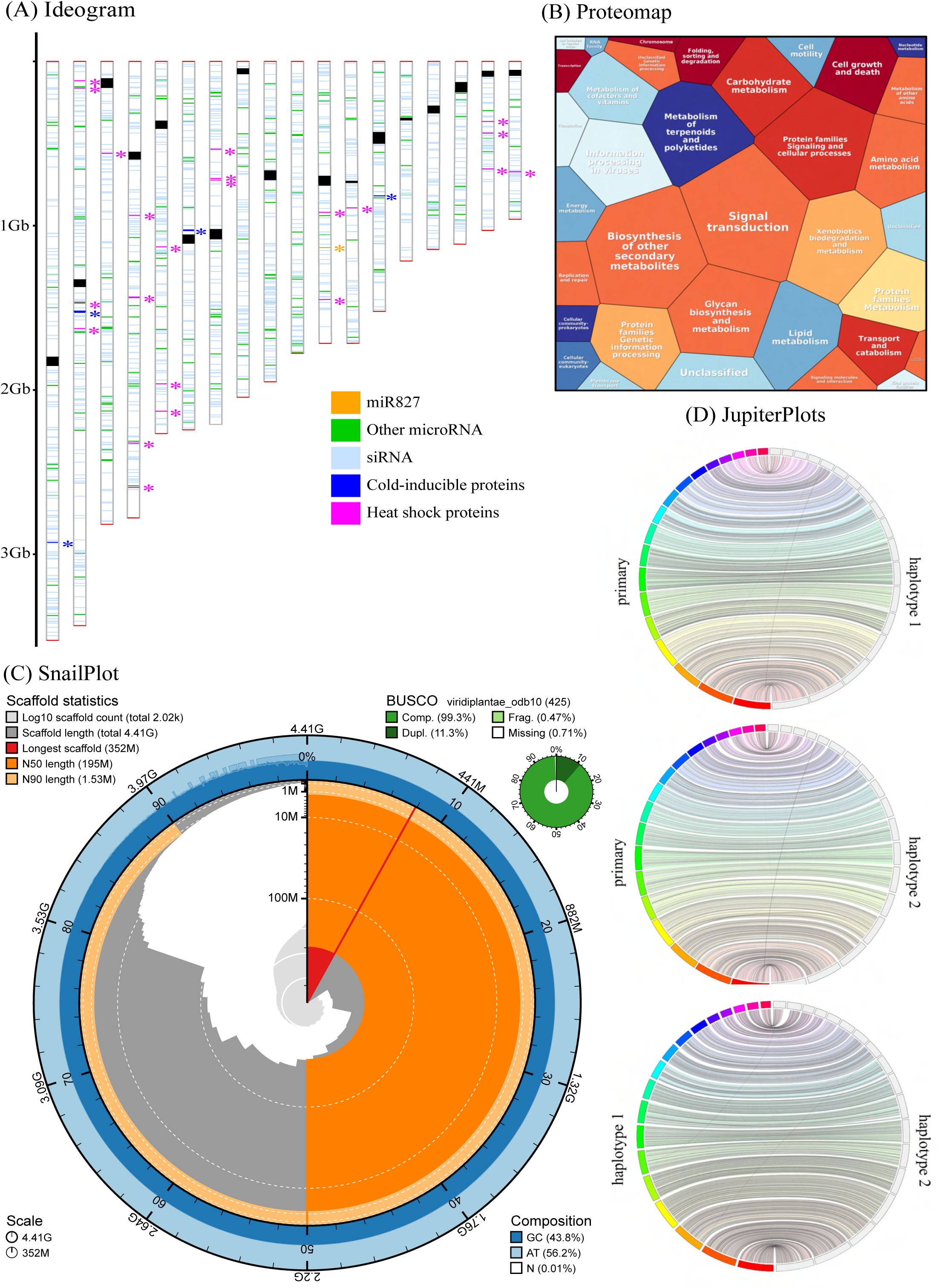
(A) ideogram showing the telomeres (red), centromeres (black), the microRNA annotated (orange, light blue, and green), and the temperature-stress proteins annotated (blue and pink). Chromosomes in the ideogram were sorted from the longest (chr 1) to the shortest one (chr 18). (B) Proteomap displaying the KEGG functional annotation groups. (C) SnailPlot showing the primary assembly statistics. (D) JupiterPlots showing the synteny between the primary and each haplotype assemblies, and between haplotypes.

The JupiterPlots (Fig. 2C) showed high levels of synteny between the primary and haplotype 1 assemblies, and between the primary and haplotype 2 assemblies, as expected given the identical chromosome numbers. However, the synteny between haplotypes revealed that scaffolds 17 and 18 are incomplete in haplotype 2.

Blobtools analysis indicated no evident contamination in any of the assemblies (Fig. S2 and Fig. S3).

### Transcriptome assembly and quality assessment

We obtained 11 Gbp of paired-end mRNA raw data, of which 97.7% passed length and quality filtering, and 1.2 Gbp of microRNA data, of which 99.8% passed length and quality filtering. The transcripts obtained from the RNA sequencing exhibited 91.8% BUSCO completeness (Table S5).

The annotation of microRNAs identified 13 known miRNAs, 47 putative miRNAs, and 4 different siRNAs (Table S6). The miRNA identified were miR156, miR159, miR160, miR164, miR166, miR167, miR168, miR172, miR395, miR396, miR528, miR535, and miR5179, and the siRNA were siRNA21, siRNA22, siRNA23, and siRNA24.

### Nuclear genome annotation

#### Repeat annotation

Genome masking revealed that 88% of the genome assemblies are repeats (Table S7). LTR retrotransposons represent most of the transposable elements in C. humilis (63%), primarily composed of two superfamilies: Try/Copia (38%) and Gypsy (25%). The remaining repeat classes, such as DNA transposons, small RNA, simple repeats, and others, accounted for less than 10% (Table 1, Table S7).

#### Gene annotation

The primary genome assembly resulted in 28,321 annotated genes. A BUSCO score of 98.3% for identified complete Viridiplantae orthologous proteins suggests high annotation completeness (C: 98.3% [S: 88.7% and D: 9.6%], F:0.7%, M:1.0%; n=425). InterProScan analysis classified 27,905 (98.5%) of the predicted proteins, of which 21,386 (76%) have a GO term associated. Finally, 24,116 (85%) of the proteins were identified in the Swiss-Prot database.

MCScanX provided the following classification of the duplicated proteins: 3,128 were singletons, 7,169 were dispersed, 1,709 were proximal, 3,986 were tandem, and 12,330 were whole-genome or segmental duplications. These proteins were clustered into 4,017 collinear groups (Fig. S4).

### Organelle genome assembly and annotation

#### Mitochondrial genome assembly

We obtained the first circular mitochondrial genome for the species C. humilis, which had a total length of 522.6 Kbp, a GC content of 45.71%, and a total of 53 annotated genes (Table S8, Fig. S5, Fig. S6). The genes obtained using MitoHifi included Complex I (nad9), Complex IV (cox1, cox3), Complex V (atp1, atp4, atp6, atp8, atp9), Cytochrome c biogenesis (ccmFc, ccmFn), Ribosome large subunit (rpl16, rpl2, rpl5), Ribosome small subunit (rps1, rps2, rps3, rps4, rps7, rps11, rps12, rps14, rps19), Intron maturase (matR), SecY-independent transporter (mttB), tRNA genes (tRNA-Asp, tRNA-Cys, tRNA-Gln, tRNA-Lys, tRNA-Met, tRNA-Phe, tRNA-Pro, tRNA-Ser, tRNA-Tyr), and other genes (CYTB, ND2, ND4, ND4L, ND5, ND6, orf118, orf142, orf192). All palm mitochondrial genomes had similar GC contents but showed variable numbers of genes within and among species (Table S9).

#### Chloroplast genome assembly

We obtained a chloroplast genome with a total length of 174.5 Kbp, a depth coverage of 44x and a GC content of 37.24% (Fig. S6 and Table S10). It has a quadripartite structure of large (LSC, 86303 bp) and small single-copy (SSC, 17925 bp) regions separated by a pair of inverted repeats (IRs, 27 bp). The genome contains 116 unique genes according to the Geseq annotation: 80 protein-coding genes, 4 ribosomal RNA genes and 32 tRNA genes (Table S10). Comparisons with the other complete chloroplast genomes of C. humilis published in NCBI showed that our obtained chloroplast genome length is higher but had similar GC content (Table S11). Moreover, structural features and number of genes of C. humilis chloroplasts are similar to those of most other Arecaceae species (Chen et al. 2022).

#### Comparison to other published palm genomes

Compared to other chromosome-level genome assemblies of palms available on NCBI, the genome of *C. humilis* has the highest values in terms of assembly length, contig and scaffold N50, and BUSCO completeness (Table S1).

In light of the observations of Giovino et al. (2024) regarding the potential adaptation of *C. humilis* to cold environments, we examined the total number of proteins involved in temperature regulation among palms. Our findings indicate that cold-inducible proteins are more prevalent in *C. humilis* than in the other tropical palms analyzed (Fig. 3).

**Fig. 3:**
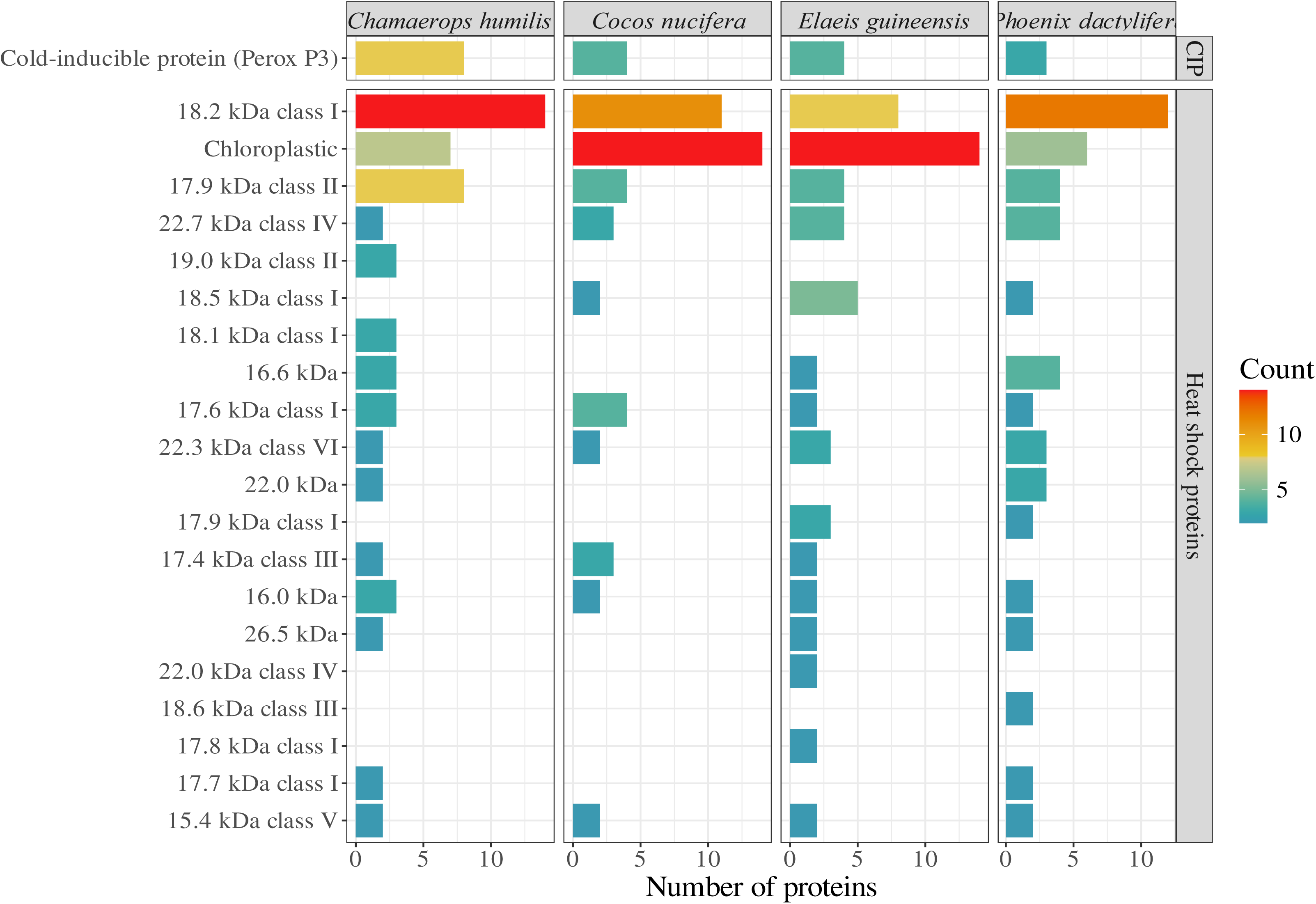
Comparison of the number of proteins involved in temperature regulation and stress response processes. The comparisons were made among *Chamaerops humilis* and three other palm species with publicly available proteomes: *Cocos nucifera*, *Elaeis guineensis*, and *Phoenix dactylifera*.

According to the phylogenomic BUSCO tree reconstruction, all the genomes from the same species were monophyletic with high clade support (bootstrap support of 1, Fig. S7). The species *C. humilis* (18 chromosomes) is sister to the genus *Phoenix* (18-19 chromosomes), which includes two sequenced species, *P. dactylifer*a and *P. roebelenii*. There are two other clades: one comprising the genera *Calamus* and *Metroxylon*, and another comprising the genera *Areca*, *Elaeis* and *Cocos* (16 chromosomes). The phylogenetic reconstruction showed a strong phylogenetic signal for chromosome number, but there was no correlation between the assembly length and chromosome number, as already suggested by Barret et al. (2019) (Table S1, Fig. S7). The obtained phylogeny is identical to that inferred by Li et al. (2021), Chen et al. (2022), and Bacon and Hill (2024).

Within angiosperms the proportion of LTR elements contribute the most to genome size variation (Wang et al. 2021). In our study, we confirm that this is also true for palms, where small genomes such as those from *Phoenix* or *Metroxylon* have lower proportions of LTR elements than large ones like those from *Chamaerops* or *Areca* (Table S1, Fig. 4). The linear regression shows an adjusted R^2^ of 0.74 (p-value = 1.08e^-5^), with a residual standard error of 0.54. The repeat divergence landscapes of the palm species showed that *C. humilis* has a higher proportion of repetitive elements at low Kimura values compared to the other palms, which instead exhibit a Gaussian distribution (Fig. 4, Fig. S8). This indicates a more recent LTR repeat activity in *C. humilis*.

**Fig. 4:**
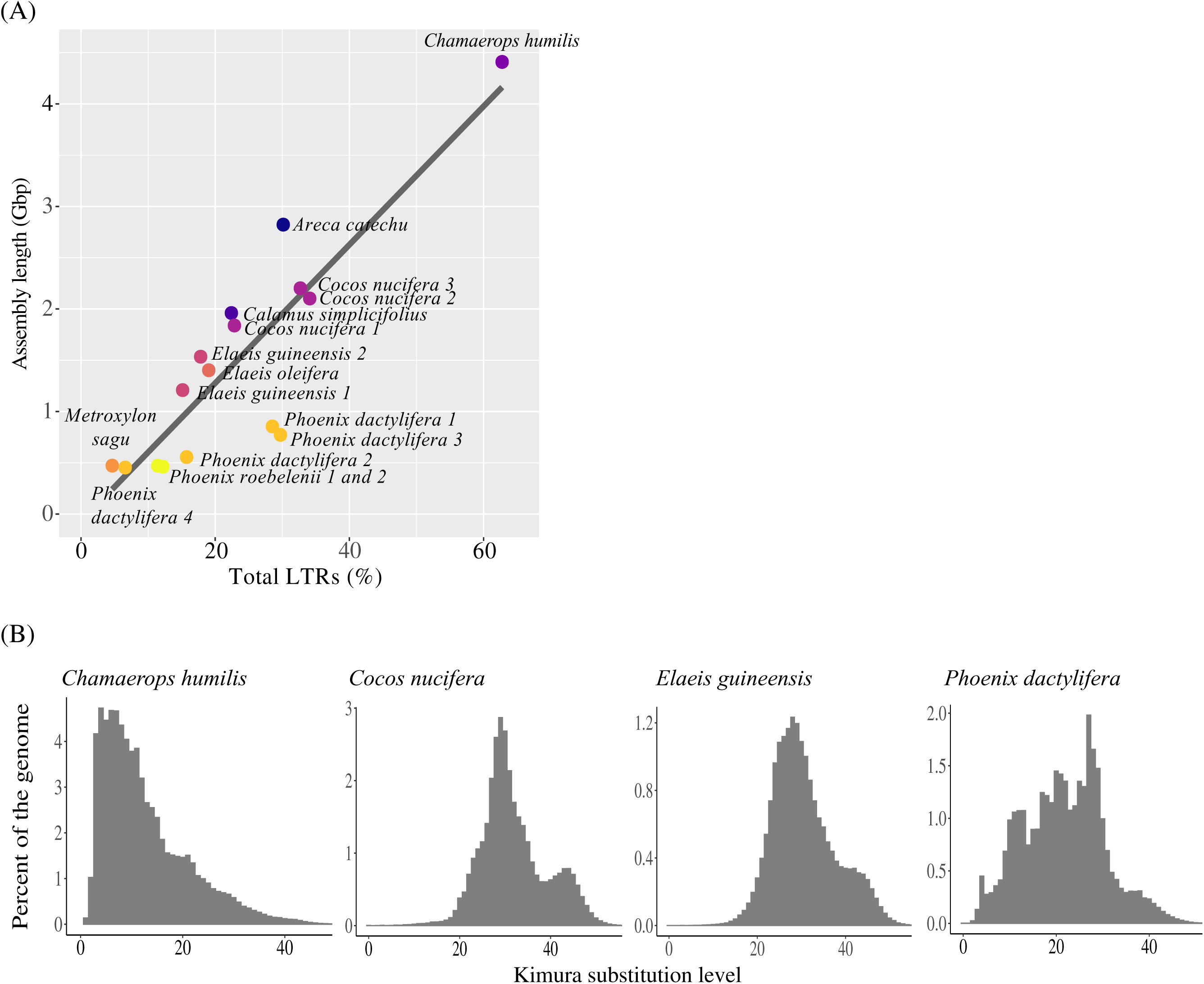
(A) Linear regression between assembly length and proportion of LTR elements present in the genome. (B) Repeat divergence landscapes of LTR elements in the main crops of the family Arecaceae .

#### Geographical origin of Goethe’s Palm

Among the ten microsatellite loci described in Giovino et al. (2023), only eight were found in the Goethe’s Palm (locus 15, locus 16, locus 19, locus 23, locus 25, locus 27, locus 37 and locus 44). Locus 25 was only present in haplotype 1, while locus 37 was only present in haplotype 2. The DAPC explained 77% of the dataset’s variability and revealed a strong geographical pattern, with axis 1 distinguishing between the western and eastern Mediterranean populations (Fig. 5).

**Fig. 5:**
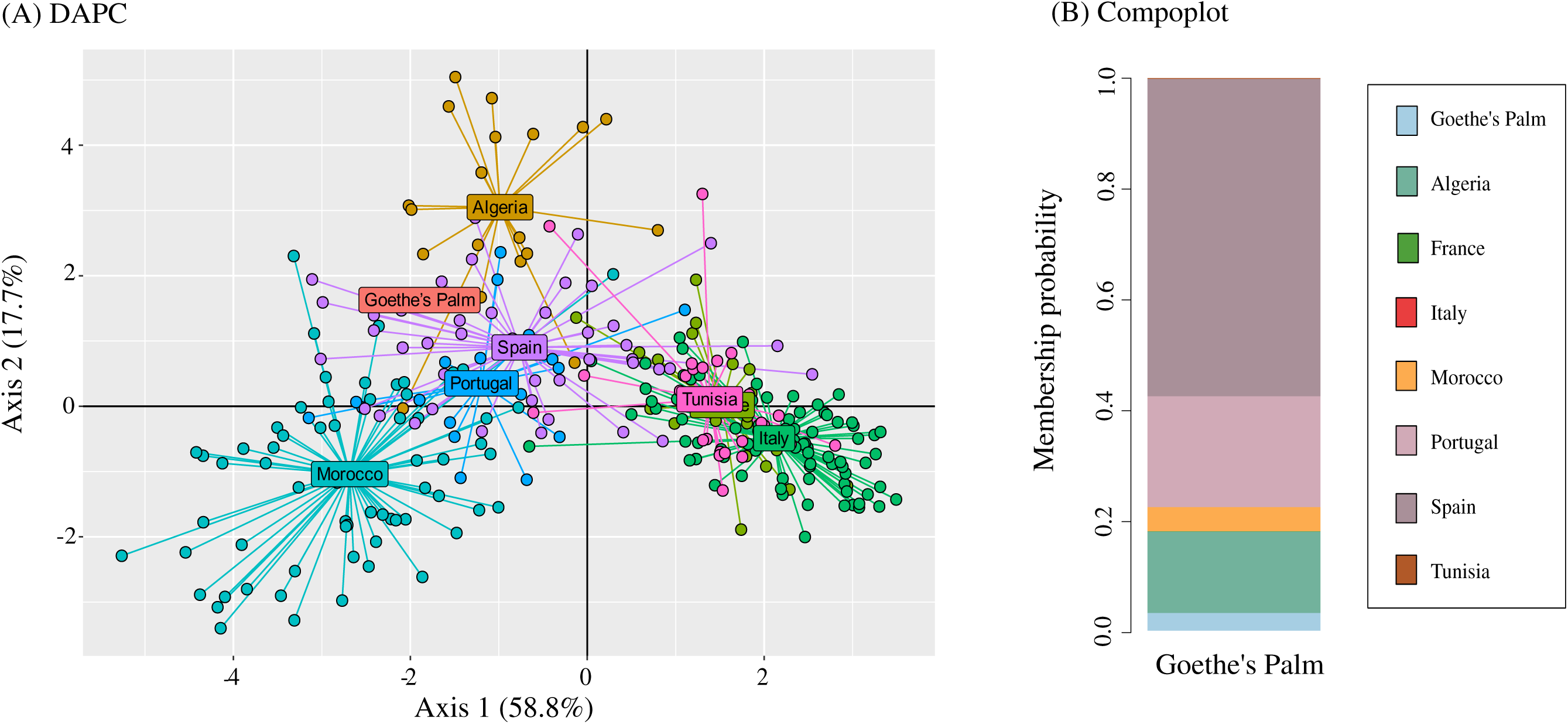
(A) DAPC showing population structure in Mediterranean dwarf palms and the potential geographic origin of Goethe’s Palm. All samples analyzed represented by filled circles are colored according to the country to which they belong. (B) Compoplot showing the most likely assignment of Goethe’s Palm to the evaluated countries. Both analyses indicate that Goethe’s Palm belongs to the Western genetic lineage of *C. humilis*, which includes populations found predominantly in Spain and Portugal.

## Discussion

The sequencing of Goethe’s Palm, the oldest cultivated palm, has yielded the most contiguous and comprehensive palm genome to date. These findings not only enhance our understanding of *Chamaerops humilis* at a molecular level but also provide valuable insights into the historical origins and evolutionary history of this specimen, shedding light on its provenance and the biogeographic lineage of the species.

### The largest and most repetitive genome in palms

The most striking feature of the *C. humilis* genome is its assembly length, making it the longest palm genome sequenced to date. However, it is important to distinguish between genome size and assembly length, as the latter may be underestimated in genomes with high proportions of repetitive elements. Previously assembled palm genomes may appear shorter than their actual size due to collapsed repeat sequences in the assemblies.

In *C. humilis*, the high transposon content, dominated by LTR transposons, contributes to this large assembly size. Repetitive elements are characteristic of larger plant genomes; for example, the genome of barley (*Hordeum vulgare* ssp. *vulgare*) comprises approximately 80-90% repetitive elements (Meyers et al., 2001; Vicient et al., 2001), while in rice (*Oryza sativa* L.), they represent 37% of the 372 Mbp genome (McCarthy et al., 2002).

Small RNAs are directly involved in gene silencing at the post-transcriptional level by inhibiting translation or by promoting sequence-specific degradation of mRNAs, with small interfering RNAs (siRNAs) and microRNAs (miRNAs) being the most important (da Silva et al. 2016). *C. humilis* genome features a diverse array of microRNAs, all of them found in other palms as well (da Silva et al., 2016; Salgado et al., 2022; Ooi et al., 2023). While the functions of many of these miRNAs in plants remain largely unknown, some have been associated with specific physiological roles. For instance, miR169 and miR408 are involved in drought resistance (Salgado et al., 2022), miR528 in salt resistance (da Silva et al., 2016), and miR160, miR164, and miR396 in root development (da Silva et al., 2016). Notably, miR528 is specific to monocot plants (Chen et al., 2019) and is present in all palms sequenced. Our study is the first to identify the miRNA family miR827 in palms, which plays a crucial role in nutrient regulation, particularly phosphate homeostasis. miR827 targets SPX domain-containing proteins involved in phosphate sensing and signaling, aiding in phosphate uptake and distribution under varying environmental conditions (e.g., Lin et al., 2013). The high content of these miRNAs and repetitive elements in *C. humilis* underscores the complexity and adaptability of its genome. Repetitive elements, such as transposons, have been shown in fact to contribute to adaptability by promoting genome plasticity, enabling rapid responses to environmental changes, and driving evolutionary innovation (Feschotte, 2008; Rebollo et al., 2010; Casacuberta and González, 2013).

### TE landscape diversity in palms suggests rapid LTR expansion bursts in C. humilis

The *C. humilis* genome revealed a high proportion of abundant TE sequences with low sequence divergence, especially of the LTR family, indicating recent TE activity. This observation aligns with the burst model of TE activity (Charlesworth & Charlesworth, 1983), which suggests that TEs proliferate in high copy rate bursts following periods of inactivity, often triggered by environmental stresses (Miousse et al., 2015; Galindo-González et al., 2017). Environmental conditions such as drought, temperature extremes, and pathogen attacks can activate dormant LTR retrotransposons, leading to their replication and reinsertion into new genomic locations (Alzohairy et al., 2014). This process increases genetic variability, potentially resulting in advantageous mutations that aid adaptation to adverse conditions (Grandbastien, 2014). Additionally, environmental stresses can alter the epigenetic landscape, influencing LTR retrotransposon activity through changes in DNA methylation and histone modifications (Srikant and Drost, 2020).

### Goethe’s Palm and the Western genetic lineage of C. humilis

The genome sequencing of Goethe’s Palm has provided new insights into its genetic lineage and origins. Our analysis indicates that Goethe’s Palm belongs to the Western genetic lineage of *C. humilis*, which includes populations found predominantly in Spain and Portugal (Giovino et al. 2023). The signal we identified in the microsatellite data suggests that the specimen, planted in 1585, likely originated from regions involved in extensive trade with Venice (Italy) during the late 16th century.

The identification of Goethe’s Palm as part of the Western lineage sheds light on the cultural and historical context of its arrival in Padua. During the Renaissance, Venice was a major trading hub, with active exchanges of goods, plants, and knowledge between Europe and North Africa. The Botanical Garden of Padua, established in 1545, benefited from these exchanges, acquiring exotic species for study and cultivation. The provenance of Goethe’s Palm can likely be traced back to these trade routes, emphasizing the interconnectedness of botanical exploration and commerce during this period. This information not only enriches our understanding of the University of Padua Botanical Garden’s collection but also highlights the significance of plant trade in shaping the diversity of plant species in European gardens.

An intriguing avenue for future research involves a comparative analysis between the genetic makeup of Goethe’s Palm and the specimen collected by Johann Wolfgang von Goethe during his visit to Padua in 1786, which is stored at the Weimar Goethe’s herbarium. Such a herbariomic study could provide valuable insights into the mutation rates and genetic changes that have occurred over the past two centuries.

### Implications for conservation and cultivation

The socio-economic importance of *C. humilis* has been recognized for decades, providing valuable resources such as fiber, food, and ornamental items. While *C. humilis* is not as commercially important as other palms, like coconut or oil palm, its role in Mediterranean countries, especially under the pressures of global change, makes it a valuable crop candidate. As a crop, defined as any cultivated plant, especially those that have been or are being domesticated (Hufford et al. 2019), *C. humilis* contributes to the livelihoods of local communities through its diverse uses.

The adaptability of *C. humilis*, particularly its wide temperature tolerance, makes it resilient to environmental fluctuations. Preliminary findings in the response of *C. humilis* to cold suggest that this species may produce a higher number of cold-induced proteins compared with tropical palm species. This resilience points out a potential role of habitat in modulating cold stress responses in palms, that may serve as a starting point for better understanding stress tolerance mechanisms in other, more economically important palms.

*Chamaerops humilis* is already endangered in parts of its natural range due to habitat loss, over-exploitation, and climate change. Botanical gardens play a crucial role in maintaining the genetic resources of species like *C. humilis* through in situ preservation and gene banks of both natural and cultivated populations. These efforts are vital for safeguarding the species’ survival and supporting future breeding initiatives, ensuring its resilience and adaptability to environmental and anthropogenic challenges.

## Conclusions

In this study, we present the most contiguous and complete palm genome to date by sequencing Goethe’s Palm, the oldest cultivated palm in the world. The high proportion of repetitive elements, particularly LTR, and the identification of key miRNAs such as miR827 provide a basis for further research into the genomic factors that contribute to the evolutionary history and ecological success of this species. This new genomic resource paves the way for more detailed investigations into the genomic diversity of *C. humilis*, offering critical insights that will significantly inform and enhance future conservation strategies.

## Supplementary Material

**Supplementary Table S1:** Summary of the main characteristics of all palm genomes published on NCBI.

**Supplementary Table S2:** Statistics for the primary assembly.

**Supplementary Table S3:** Statistics for the assembly of haplotype 1.

**Supplementary Table S4:** Statistics for the assembly of haplotype 2.

**Supplementary Table S5:** Total number of transcripts and BUSCO completeness of the RNA data.

**Supplementary Table S6:** microRNA identification.

**Supplementary Table S7:** Repeat masking table.

**Supplementary Table S8:** Statistics for the *C. humilis* mitochondrial genome.

**Supplementary Table S9:** Statistics for all mitochondrion assemblies available in NCBI for the family *Arecaceae*.

**Supplementary Table S10:** Plastid genome assembly statistics for *C. humilis*.

**Supplementary Table S11:** Statistics for all plastid assemblies available in NCBI for the genus *Chamaerops*.

**Supplementary Fig. S1:** Hi-C contact density map.

**Supplementary Fig. S2:** BlobPlots.

**Supplementary Fig. S3:** ReadCovPlots.

**Supplementary Fig. S4:** Synteny visualization of the collinearity in the predicted proteins of the primary assembly.

**Supplementary Fig. S5:** Mitochondrial annotation obtained with MitoHifi for the primary genome assembly.

**Supplementary Fig. S6:** Organelles genome annotation.

**Supplementary Fig. S7:** Phylogenomic BUSCO tree of all published genomes.

**Supplementary Fig. S8:** Repeat divergence landscapes in several species of the family Arecaceae.

## Acknowledgments

We thank the LOEWE Centre for Translational Biodiversity Genomics (LOEWE-TBG), especially Alexander Ben Hamadou and Charlotte Gerheim, for their wet lab support. We acknowledge the Genome Technology Center (RGTC) at Radboudumc for access to the Sequencing Core Facility (Nijmegen, Netherlands), where PacBio SMRT sequencing was performed on the Sequel IIe platform. We are grateful to Rhonda Struminger (Botanical Garden of Padua) for her comments on the manuscript and assistance with English language review. We also thank Carlo Calore and the Ufficio Eventi Permanenti (Botanical Garden of Padua) for providing palm images and the garden map.

## Data availability

All data have been submitted to the NCBI BioSample accession number SAMN39481424. The GenBank assembly numbers are as follows: the GCA_042465325.1 has been designated for haplotype 1, GCA_042465335.1 for haplotype 2, and GCA_042465385.1 for the primary genome assembly. The BioProject accession numbers are as follows: PRJNA1066108 has been assigned to haplotype 1, PRJNA1066109 to haplotype 2, and PRJNA1066107 to the primary genome assembly. The RNA sequencing data have been submitted to the NCBI BioProject accession number PRJNA1066107. The accession numbers SRR29081397 and SRR29081396 pertain to mRNA and microRNA, respectively. The mitochondrial and plastid assemblies have been uploaded to the Zenodo repository and can be accessed via the following DOI number: https://doi.org/10.5281/zenodo.14872859.

