## supplementary material for "Sequencing a botanical monument: a chromosome-level assembly of the 400-year-old Goethe’s Palm (*Chamaerops humilis* L.) at the Botanical Garden of the University of Padua (Italy)"

**Table S1:** Summary of the main characteristics of all palm genomes published on NCBI. The genome statistics (genome length, N50, and GC content) were calculated with Quast, and the percent of complete BUSCO genes was calculated with BUSCO. The LTR elements were calculated with RepeatMasker. The other information was extracted directly from the NCBI website.

| NCBI accession number | Species | Sequencing Technology | Assembly level | Genome length (Gbp) | N50 (bp) | GC (%) | Genome coverage | Complete BUSCO genes (%) | Annotation | LTR (%) |
|---|---|---|---|---|---|---|---|---|---|---|
| GCA_021397845.1 | *Areca catechu* | PacBio | Chromosome (16 Chr) | 2.82 | 186523490 | 41.45 | 100x | 95 | Not performed | 30.14 |
| GCA_900491605.1 | *Calamus simplicifolius* | Missing data | Scaffold | 1.96 | 803014 | 41.06 | 270x | 91 | Not performed | 22.41 |
| GCA_003604295.1 | *Cocos nucifera 1* | Illumina; PacBio | Scaffold | 1.84 | 85564 | 37.31 | 53x | 93 | Not performed | 22.86 |
| GCA_006176705.1 | *Cocos nucifera 2* | Illumina MiSeq; PacBio RSII; HiRise pipeline | Scaffold | 2.10 | 570487 | 37.73 | 50x | 95 | Not performed | 34.09 |
| GCA_008124465.1 | *Cocos nucifera 3* | Illumina HiSeq2000 | Chromosome (16 Chr) | 2.20 | 1217559 | 37.30 | 173x | 90 | Performed | 32.69 |
| GCA_015461965.1 | *Elaeis guineensis 1* | Linkage mapping | Chromosome (16 Chr) | 1.21 | 76242894 | 37.05 | 16x | 93 | Not performed | 15.14 |
| GCF_000442705.1 | *Elaeis guineensis 2* | 454 | Chromosome (16 Chr) | 1.54 | 1268079 | 37.21 | 16x | 94 | Performed | 17.83 |
| GCA_000441515.1 | *Elaeis oleifera* | 454 | Scaffold | 1.40 | 333109 | 37.99 | 16x | 75 | Not performed | 19.04 |
| GCA_017589505.3 | *Metroxylon sagu* | Oxford Nanopore; Illumina NovaSeq | Scaffold | 0.47 | 1534258 | 36.08 | 50x | 95 | Not performed | 4.65 |
| GCA_000181215.3 | *Phoenix dactylifera 1* | PacBio RSII; Illumina HiSeq; 10X Genomics | Chromosome (18 Chr) | 0.85 | 51272265 | 40.37 | 120x | 92 | Not performed | 28.52 |
| GCF_000413155.1 | *Phoenix dactylifera 2* | 454; SOLiD; | Scaffold | 0.56 | 335289 | 39.33 | 139x | 90 | Performed | 19.04 |

| NCBI accession number | Species | Sequencing Technology | Assembly level | Genome length (Gbp) | N50 (bp) | GC (%) | Genome coverage | Complete BUSCO genes (%) | Annotation | LTR (%) |
|---|---|---|---|---|---|---|---|---|---|---|
| | | ABI3730 | | | | | | | | |
| GCF_009389715.1 | *Phoenix dactylifera 3* | PacBio | Chromosome (19 Chr) | 0.77 | 4728343 | 40.29 | 100x | 95 | Performed | 29.72 |
| GCA_025769935.1 | *Phoenix roebelenii 1* | Illumina | Scaffold | 0.47 | 10503348 | 39.74 | 95x | 93 | Not performed | 11.43 |
| GCA_028774785.1 | *Phoenix roebelenii 2* | Illumina HiSeq | Scaffold | 0.46 | 569782 | 39.37 | 50x | 83 | Not performed | 12.20 |
| GCA_007821505.1 | *Phoenix dactylifera 4* | Illumina HiSeq | Scaffold | 0.45 | 5673 | 38.90 | 154x | 55 | Not performed | 6.59 |

**Table S2:** Statistics for the primary genome assembly calculated with Quast at scaffold level, and with Assemblathon2 at contig level. In the 'Hi-C pipeline first run' and 'Hi-C pipeline second run' columns, BUSCO completeness was calculated after running TGS-Gap Closer.

| | Statistics for the primary assembly | | | | | |
|---|---|---|---|---|---|---|
| | Hifiasm | Hi-C pipeline first run | | Hi-C pipeline second run | | Assembly without contamination and organelles |
| | | Pre Gap closer | Post Gap closer | Pre Gap closer | Post Gap closer | |
| **SCAFFOLDS:** | | | | | | |
| Scaffolds (>= 0 bp) | - | 2317 | 2317 | 2015 | 2015 | 1986 |
| Scaffolds (>= 50000 bp) | - | 1828 | 1828 | 1536 | 1536 | 1524 |
| Largest scaffold (bp) | - | 338985879 | 339009363 | 352386025 | 352408672 | 352408672 |
| Total length (>= 0 bp) | - | 4407895112 | 4408031063 | 4407957912 | 4408125824 | 4406632840 |
| Total length (>= 50000 bp) | - | 4393734959 | 4393870910 | 4394124192 | 4394292104 | 4393395764 |
| GC (%) | - | 43.80 | 43.80 | 43.80 | 43.80 | 43.80 |
| N50 | - | 195019846 | 195022640 | 195019846 | 195022367 | 195022367 |
| N90 | - | 1112000 | 1112000 | 1527355 | 1527355 | 1531061 |
| L50 | - | 9 | 9 | 9 | 9 | 9 |
| L90 | - | 197 | 197 | 134 | 134 | 133 |
| N's per 100 kbp | - | 5.20 | 4.71 | 6.63 | 5.93 | 5.94 |
| **CONTIGS:** | | | | | | |
| Number of contigs | 2897 | 3464 | 3356 | 3476 | 3323 | 3294 |
| Number of contigs in scaffolds | 0 | 1225 | 1109 | 1687 | 1501 | 1501 |
| Total length of contigs | 4407665712 | 4407665712 | 4407823263 | 4407665712 | 4407864224 | 4406371240 |
| Largest contig (bp) | 28621313 | 28621313 | 28621313 | 28621313 | 28621313 | 28621313 |

| | Statistics for the primary assembly | | | | | |
|---|---|---|---|---|---|---|
| | **Hifiasm** | **Hi-C pipeline first run** | | **Hi-C pipeline second run** | | **Assembly without contamination and organelles** |
| | | **Pre Gap closer** | **Post Gap closer** | **Pre Gap closer** | **Post Gap closer** | |
| Median contig size (bp) | 476405 | 353000 | 327696 | 352239 | 339334 | 348000 |
| Shortest contig (bp) | 6241 | 1000 | 1000 | 950 | 950 | 950 |
| N50 | 4424378 | 3866000 | 4366744 | 3866000 | 4380000 | 4383116 |
| L50 | 280 | 320 | 288 | 320 | 287 | 286 |
| **BUSCO completeness (n = 425):** | | | | | | |
| Busco genes for all scaffolds and contigs | C: 99.3% [S: 87.1% and D: 12.2%], F: 0.5%, M: 0.2% | C: 99.3% [S: 87.8% and D: 11.5%], F: 0.5%, M: 0.2% | | C: 99.3% [S: 88.0% and D: 11.3%], F: 0.5%, M: 0.2% | | C: 99.3% [S: 88.0% and D: 11.3%], F: 0.5%, M: 0.2% |

**Table S3:** Statistics for the assembly of haplotype 1 calculated with Quast at scaffold level, and with Assemblathon2 at contig level. In the 'Hi-C pipeline first run' and 'Hi-C pipeline second run' columns, BUSCO completeness was calculated after running TGS-Gap Closer.

| | Statistics for the Haplotype 1 assembly | | | | | |
|---|---|---|---|---|---|---|
| | Hifiasm | Hi-C pipeline first run | | Hi-C pipeline second run | | Assembly without contamination and organelles |
| | | Pre Gap closer | Post Gap closer | Pre Gap closer | Post Gap closer | |
| **SCAFFOLDS:** | | | | | | |
| Scaffolds (>= 0 bp) | - | 1861 | 1861 | 1818 | 1818 | 1783 |
| Scaffolds (>= 50000 bp) | - | 1099 | 1099 | 1062 | 1062 | 1050 |
| Largest scaffold (bp) | - | 342648678 | 342751203 | 353581407 | 353674888 | 353674888 |
| Total length (>= 0 bp) | - | 4040616072 | 4041394661 | 4040626672 | 4041312864 | 4039555345 |
| Total length (>= 50000 bp) | - | 4020898855 | 4021677444 | 4021081271 | 4021767463 | 4020938903 |
| GC (%) | - | 43.82 | 43.82 | 43.82 | 43.82 | 43.82 |
| N50 | - | 194726223 | 194771343 | 211806283 | 211881467 | 211881467 |
| N90 | - | 3516590 | 3516900 | 4028457 | 4028457 | 4051748 |
| L50 | - | 8 | 8 | 8 | 8 | 8 |
| L90 | - | 43 | 43 | 30 | 30 | 29 |
| N's per 100 kbp | - | 18.22 | 15.73 | 18.49 | 16.07 | 16.08 |
| **CONTIGS:** | | | | | | |
| Number of contigs | 5193 | 5543 | 5040 | 5553 | 5065 | 5030 |
| Number of contigs in scaffolds | 0 | 3823 | 3309 | 3873 | 3366 | 3366 |
| Total length of contigs | 4039879672 | 4039879672 | 4040758861 | 4039879672 | 4040663464 | 4038905945 |
| Largest contig (bp) | 21541770 | 21541770 | 21541770 | 21541770 | 21541770 | 21541770 |

| | Statistics for the Haplotype 1 assembly | | | | | |
|---|---|---|---|---|---|---|
| | Hifiasm | Hi-C pipeline first run | | Hi-C pipeline second run | | Assembly without contamination and organelles |
| | | Pre Gap closer | Post Gap closer | Pre Gap closer | Post Gap closer | |
| Median contig size (bp) | 352136 | 312658 | 339207 | 312312 | 334656 | 341240 |
| Shortest contig (bp) | 3082 | 1000 | 1000 | 863 | 863 | 863 |
| N50 | 1796000 | 1759376 | 1963804 | 1759376 | 1982289 | 1982289 |
| L50 | 555 | 588 | 530 | 588 | 527 | 527 |
| **BUSCO completeness (n = 425):** | | | | | | |
| Busco genes for all scaffolds and contigs | C: 95.5% [S: 86.8% and D: 8.7%], F: 1.4%, M: 3.1% | C: 96.2% [S: 88.9% and D: 7.3%], F: 0.7%, M: 3.1% | | C: 96.0% [S: 88.7% and D: 7.3%], F: 0.9%, M: 3.1% | | C: 96.0% [S: 88.7% and D: 7.3%], F: 0.9%, M: 3.1% |

**Table S4:** Statistics for the assembly of haplotype 2 calculated with Quast at scaffold level, and with Assemblathon2 at contig level. In the 'Hi-C pipeline first run' and 'Hi-C pipeline second run' columns, BUSCO completeness was calculated after running TGS-Gap Closer.

| | **Statistics for the Haplotype 2 assembly** | | | | | |
|---|---|---|---|---|---|---|
| | **Hifiasm** | **Hi-C pipeline first run** | | **Hi-C pipeline second run** | | **Assembly without contamination and organelles** |
| | | **Pre Gap closer** | **Post Gap closer** | **Pre Gap closer** | **Post Gap closer** | |
| **SCAFFOLDS:** | | | | | | |
| Scaffolds (>= 0 bp) | - | 1489 | 1489 | 1458 | 1458 | 1432 |
| Scaffolds (>= 50000 bp) | - | 1128 | 1128 | 1097 | 1097 | 1084 |
| Largest scaffold (bp) | - | 309728793 | 309810774 | 325363200 | 325404666 | 325404666 |
| Total length (>= 0 bp) | - | 3921987479 | 3922895508 | 3921995879 | 3922899159 | 3920974641 |
| Total length (>= 50000 bp) | - | 3912001573 | 3912909602 | 3912098491 | 3913001771 | 3911604370 |
| GC (%) | - | 43.81 | 43.81 | 43.81 | 43.81 | 43.81 |
| N50 | - | 174121730 | 174169707 | 206042248 | 206085499 | 206085499 |
| N90 | - | 3718888 | 3718888 | 4270963 | 4270963 | 4430893 |
| L50 | - | 9 | 9 | 8 | 8 | 8 |
| L90 | - | 42 | 42 | 32 | 32 | 31 |
| N's per 100 kbp | - | 18.82 | 16.11 | 19.03 | 16.33 | 16.33 |
| **CONTIGS:** | | | | | | |
| Number of contigs | 4833 | 5179 | 4648 | 5190 | 4662 | 4634 |
| Number of contigs in scaffolds | 0 | 3835 | 3292 | 3873 | 3334 | 3331 |
| Total length of contigs | 3921249479 | 3921249479 | 3922263708 | 3921249479 | 3922258359 | 3920334241 |
| Largest contig (bp) | 27303349 | 20663968 | 20663968 | 20663968 | 20663968 | 20663968 |

| | Statistics for the Haplotype 2 assembly | | | | | |
|---|---|---|---|---|---|---|
| | Hifiasm | Hi-C pipeline first run | | Hi-C pipeline second run | | Assembly without contamination and organelles |
| | | Pre Gap closer | Post Gap closer | Pre Gap closer | Post Gap closer | |
| Median contig size (bp) | 405490 | 360740 | 401260 | 359513 | 401000 | 407000 |
| Shortest contig (bp) | 8230 | 1000 | 1000 | 1000 | 1000 | 1000 |
| N50 | 1675054 | 1622158 | 1833067 | 1622158 | 1818675 | 1818675 |
| L50 | 592 | 622 | 556 | 622 | 560 | 560 |
| **BUSCO completeness (n = 425):** | | | | | | |
| Busco genes for all scaffolds and contigs | C: 96.4% [S: 88.2% and D: 8.2%], F: 0.7%, M: 2.9% | C: 96.5% [S: 88.0% and D: 8.5%], F: 0.7%, M: 2.8% | | C: 96.5% [S: 88.0% and D: 8.5%], F: 0.7%, M: 2.8% | | C: 96.5% [S: 88.0% and D: 8.5%], F: 0.7%, M: 2.8% |

**Table S5:** Total number of transcripts and BUSCO completeness of the mRNA sequenced using as a reference genome the primary genome assembly.

| | **mRNA** |
|---|---|
| Number of transcripts | 76,940 |
| BUSCO completeness (n: 425) | C: 91.8% [S: 39.3% and D: 52.5%], F: 6.4%, M: 1.8% |

**Table S6:** microRNA identification using Arecaceae family from sRNAanno reference dataset.

| **Type of microRNA** | **Number of microRNA found** | **microRNA identification (abundance of the sRNA found)** |
|---|---|---|
| miRNA | 13 | **Identification confirmed:** miR156 (1), miR159 (2), miR162 (1), miR164 (4), miR166 (2), miR167 (1), miR171 (2), miR390 (1), miR396 (5), miR827 (2), miR5179 (1), miRN22 (2), unknown (2). |
| | 47 | **Identification unconfirmed:** miR160, miR168, miR169, miR172, miR319, miR391, miR393, miR394, miR395, miR397, miR399, miR408, miR477, miR479, miR482, miR528, miR529, miR535, miR536, miR828, miR1432, miR2118, miR3627, miR3711, miRN1, miRN2, miRN3, miRN4, miRN5, miRN6, miRN7, miRN8, miRN9, miRN10, miRN11, miRN12, miRN13, miRN14, miRN15, miRN16, miRN19, miRN20, miRN21, miRN23, unknown. |
| siRNA21 | 216 | |
| siRNA22 | 93 | |
| siRNA23 | 7 | |
| siRNA24 | 1768 | |
| Unknown | 14784 | |

**Table S7:** Repeat masking table for the primary genome assembly. The number of elements include repeats fragmented by insertions or deletions as one element.

| | **Number of elements** | **Length occupied (bp)** |
|---|---|---|
| 1. Bases masked | 3884854618 | |
| 2. Retroelements: | 2066693 | 2912216904 |
| 2.1. SINEs | 41982 | 18629799 |
| 2.2. Penelope | 3452 | 566170 |
| 2.3. LINEs: | 194598 | 125421176 |
| L2/CR1/Rex | 2919 | 588749 |
| R1/LOA/Jockey | 3915 | 1528239 |
| R2/R4/NeSL | 10060 | 4607522 |
| RTE/Bov-B | 23289 | 6598817 |
| L1/CIN4 | 95907 | 77271221 |
| 2.4. LTR elements: | 1830113 | 2768165929 |
| Ty1/Copia | 945565 | 1661956011 |
| Gypsy/DIRS1 | 872498 | 1095547454 |
| Retroviral | 141 | 168176 |
| 3. DNA transposons: | 366177 | 141502739 |
| hobo-Activator | 113650 | 43770281 |
| Tc1-IS630-Pogo | 10292 | 3634630 |
| Tourist/Harbinger | 16920 | 3666821 |
| Other (Mirage, P-element, Transib) | 7495 | 1014378 |
| 4. Rolling-circles | 13297 | 6450410 |
| 5. Unclassified | 1936256 | 682943417 |
| 6. Total interspersed repeats | - | 3736663060 |
| 7. Small RNA | 51972 | 31253510 |
| 8. Satellites | 56385 | 41043834 |
| 9. Simple repeats | 1377989 | 68001968 |
| 10. Low complexity | 306594 | 19435865 |

**Table S8:** Statistics for the *C. humilis* mitochondrial genome assembled using the primary chromosome-level genome assembly. The statistics were obtained after running MitoHifi and Quast tools.

| | **Mitochondrion assembly statistics** | |
|---|---|---|
| | **Hi-C pipeline first run** | **Hi-C pipeline second run** |
| **MitoHifi statistics** | | |
| Circular | True | True |
| Number of genes | 53 | 53 |
| Number of scaffolds of the palm genome belonging to the mitochondrion | 19 | 19 |
| **Quast statistics** | | |
| Total length (bp) | 522623 | 522623 |
| Number of contigs | 1 | 1 |
| GC content (%) | 45.71 | 45.71 |

**Table S9:** Statistics for all mitochondrion assemblies available in NCBI for the family Arecaceae. The GC content was calculated with Quast and the number of genes with GeSeq.

| NCBI accession number | Species | Reference | Sequencing technology | Circular | Total length of scaffolds (bp) | GC content (%) | Number of genes |
|---|---|---|---|---|---|---|---|
| This study | *Chamaerops humilis* | This study | PacBio | Circular | 522623 | 45.71 | • **blatN:** 41 genes, 4 rRNA, 37 tRNA.<br>• **blatX:** 37 genes. |
| PP035767 | *Cocos nucifera* | Unpublished | PacBio | Circular | 370451 | 45.45 | • **blatN:** 31 genes, 7 rRNA, 24 tRNA.<br>• **blatX:** 19 genes. |
| NC_031696 (identical to KX028885) | *Cocos nucifera* | Unpublished | Not specified | Circular | 678653 | 45.52 | • **blatN:** 54 genes, 7 rRNA, 47 tRNA.<br>• **blatX:** 70 genes. |
| JN375330 (identical to NC_016740) | *Phoenix dactylifera* | Fang et al. (2012) | Roche/454 | Circular | 715001 | 45.15 | • **blatN:** 46 genes, 3 rRNA, 43 tRNA.<br>• **blatX:** 35 genes. |
| MH176158 | *Phoenix dactylifera* | Direct submission | Illumina | Circular | 715094 | 45.12 | • **blatN:** 46 genes, 6 rRNA, 40 tRNA.<br>• **blatX:** 38 genes. |
| MG257490 (Unverified) | *Phoenix dactylifera* | Unpublished | Not specified | Linear | 585493 | 44.79 | • **blatN:** 33 genes, 3 rRNA, 30 tRNA.<br>• **blatX:**11 genes. |
| BK059358 | *Phoenix roebelenii* | Chakravartty and Reddy Neelapu (2023) | Illumina | Circular | 482735 | 46.23 | • **blatN:** 31 genes, 4 rRNA, 27 tRNA.<br>• **blatX:** 26 genes. |

**Table S10:** Plastid genome assembly statistics for *C. humilis*.

| | **Plastid assembly statistics** |
|---|---|
| **Quast statistics** | |
| Total length (bp) | 174485 |
| Largest contig (bp) | 128610 |
| Shorter contig (bp) | 45875 |
| GC (%) | 37.24 |
| N50 | 128610 |
| N's per 100 kbp | 0 |
| Number of contigs | 2 |
| **Annotation** | |
| GEseq annotation | 116 genes:<br>• **<u>80 protein-coding genes:</u> ATP synthase** (atpA, atpB, atpE, atpF, atpH, atpI), **Cytochrome b6/f** (petA, petB, petD, petG, petL, perN), **Large subunit ribosomal proteins** (rpl14, rpl16, rpl2, rpl20, rpl22, rpl23, rpl32, rpl33, rpl36), **NADH oxidoreductase** (ndhA, ndhB, ndhC, ndhD, ndhE, ndhF, ndhG, ndhH, ndhI, ndhJ, ndhK), **Photosystem I** (psaA, psaB, psaC, psaI, psaJ), **Photosystem II** (psb30, psbA, psbB, psbC, psbD, psbE, psbF, psbH, psbI, psbJ, psbK, psbL, psbM, psbT, psbZ), **RNAP** (rpoA, rpoB, rpoC1, rpoC2), **Rubisco** (rbcL), **Small subunit ribosomal proteins** (rps2, rps3, rps4, rps7, rps8, rps11, rps12, rps14, rps15, rps16, rps18, rps19), **other proteins** (accD, ccsA, cemA, clpP1, infA, matK, pafI, pafII, pbf1), **Proteins of unknown function** (ycf1, ycf2).<br>• **<u>4 ribosomal RNA genes</u>**: rrn4.5, rrn5, rrn16, rrn23.<br>• **<u>32 tRNA genes</u>**: trnA-UGC, trnC-ACA, trnC-GCA, trnD-GUC, trnE-UUC, trnF-GAA, trnfM-CAU, trnG-GCC, trnG-UCC, trnH-GUG, trnI-GAU, trnK-UUU, trnL-CAA, trnL-GAG, trnL-UAA, trnL-UAG, trnM-CAU, trnN-GUU, trnP-UGG, trnQ-UUG, trnR-ACG, trnR-UCU, trnS-CGA, trnS-GCU, trnS-GGA, trnS-UGA, trnT-GGU, trnT-UGU, trnV-GAC, trnV-UAC, trnW-CCA, trnY-GUA |

| | |
|---|---|
| PGA annotation | 102 genes:<br>• **68 protein-coding genes:** The same genes than those obtained with GEseq but without rpl2, rpl23, rpl32, ndhB. ndhE, ndhH, ndhI, psbN, psb30, rps7, rps19, ccsA, pafI, pafII, and pbf1. In addition, three more genes were found in the PGA annotation (psbN, ycf3, ycf4).<br>• **4 ribosomal RNA genes**: The same genes than those obtained with Geseq.<br>• **30 tRNA genes**: trnA-UGC, trnC-GCA, trnD-GUC, trnE-UUC, trnF-GAA, trnG-GCC, trnG-UCC, trnH-GUG, trnI-CAU, trnI-GAU, trnK-UUU, trnL-CAA, trnL-UAA, trnL-UAG, trnM-CAU, trnN-GUU, trnP-UGG, trnQ-UUG, trnR-ACG, trnR-UCU, trnS-GCU, trnS-GGA, trnS-UGA, trnT-GGU, trnT-UGU, trnV-GAC, trnV-UAC, trnW-CCA, trnY-GUA, trnfM-CAU. |
| **ptGAUL statistics** | |
| Mean coverage | 44x |

**Table S11:** Statistics calculated with Quast and PGA for all plastid assemblies available in NCBI for the genus *Chamaerops*.

| **NCBI accession number** | **Our plastid** | **ON248747** | **NC_029967** | **KT312935** |
|---|---|---|---|---|
| Reference | This study | Yao et al. (2023) | Unpublished | Unpublished |
| Sequencing technology | PacBio | Illumina | Illumina | Illumina |
| Total length of scaffolds (bp) | 174485 | 158646 | 158653 | 158653 |
| PGA annotation | 102 | 113 | 113 | 113 |
| GeSeq genes annotation | • **blatX:** 74<br>• **Chloe:** 115<br>• **HMMER:** 136 | • **blatX:** 74<br>• **Chloe:** 89<br>• **HMMER:** 131 | • **blatX:** 89<br>• **Chloe:** 89<br>• **HMMER:** 131 | • **BlatX:** 89<br>• **Chloe:** 89<br>• **HMMER:** 131 |
| GC content (%) | 37.24 | 37.19 | 37.19 | 37.19 |

**Fig. S1:** Hi-C contact density map (A) after the first run of the Arima HiC pipeline for the (A1) Primary, (A2) Haplotype 1, and (A3) Haplotype 2 assemblies; and (B) after the second run of the Arima HiC pipeline for the (B1) primary, (B2) Haplotype 1, and (B3) Haplotype 2 assemblies.

First run of the Arima Hi-C pipeline:

(A1) Primary

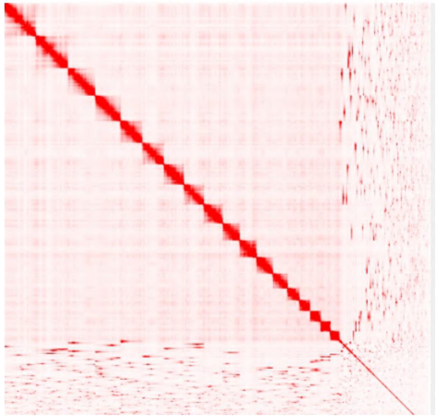

(A2) Haplotype 1

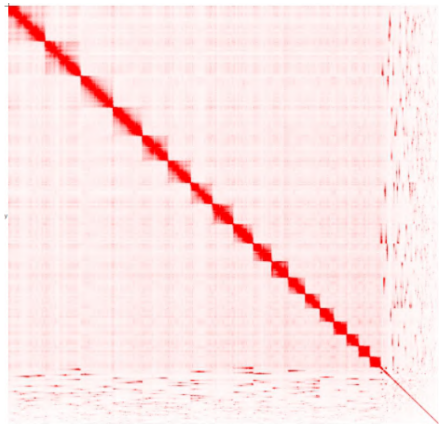

(A3) Haplotype 2

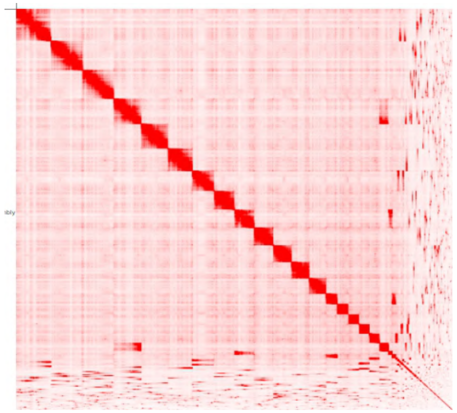

(A) Second run of the Arima Hi-C pipeline:

(B1) Primary

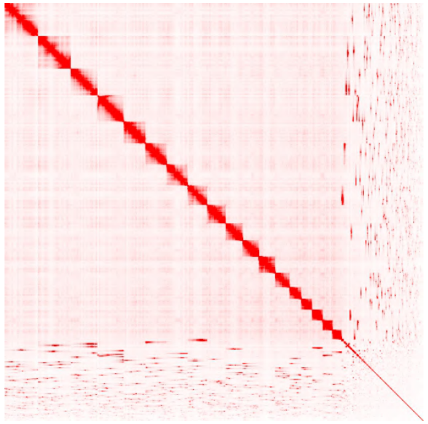

(B2) Haplotype 1

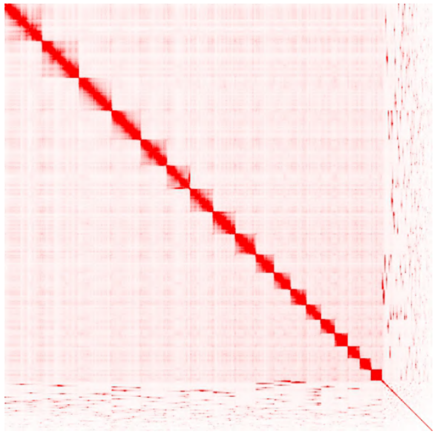

(B3) Haplotype 2

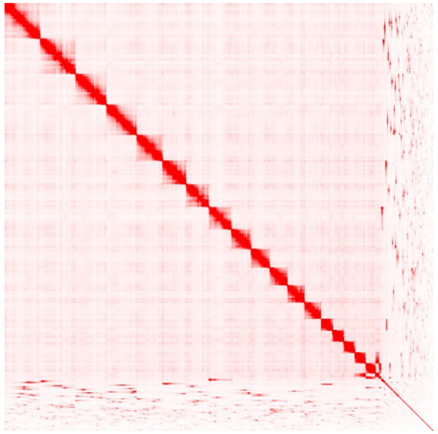

**Fig. S2:** BlobPlots comparing GC content (x axis), sequencing depth of PacBio reads (y axis), and taxonomic assignment of contigs (colors).

(A) Primary:

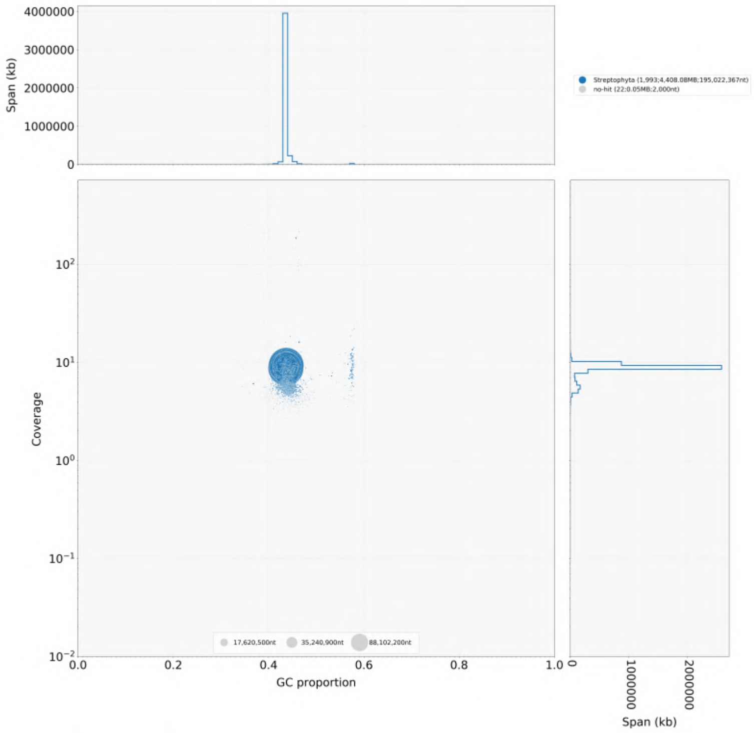


Haplotype 1:

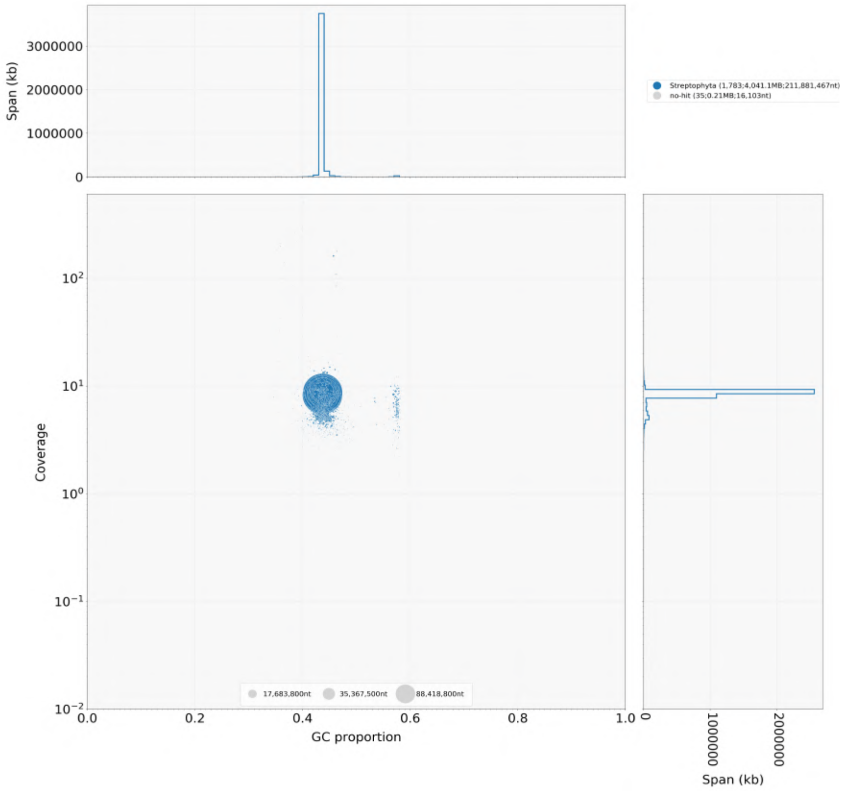

(B) Haplotype 2:

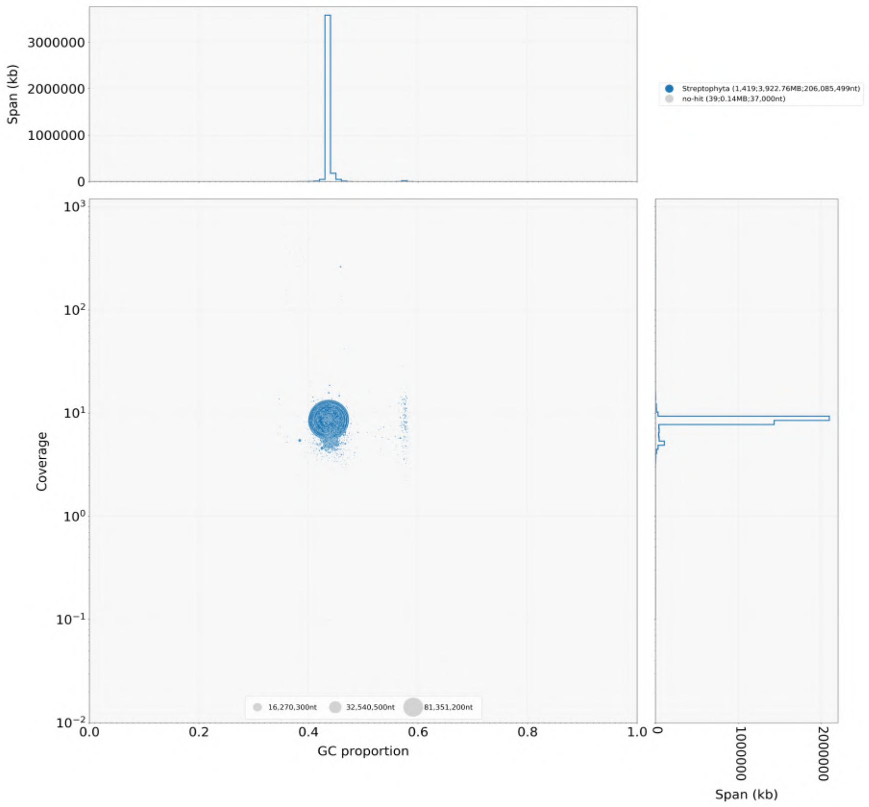


**Fig. S3:** ReadCovPlots visualising the proportion of reads of a library that are unmapped or mapped against the genome, and the percentage of mapped reads by taxonomic group for the (A) Primary, (B) Haplotype 1, and (C) Haplotype 2 obtained using Blobtools.

(A) Primary

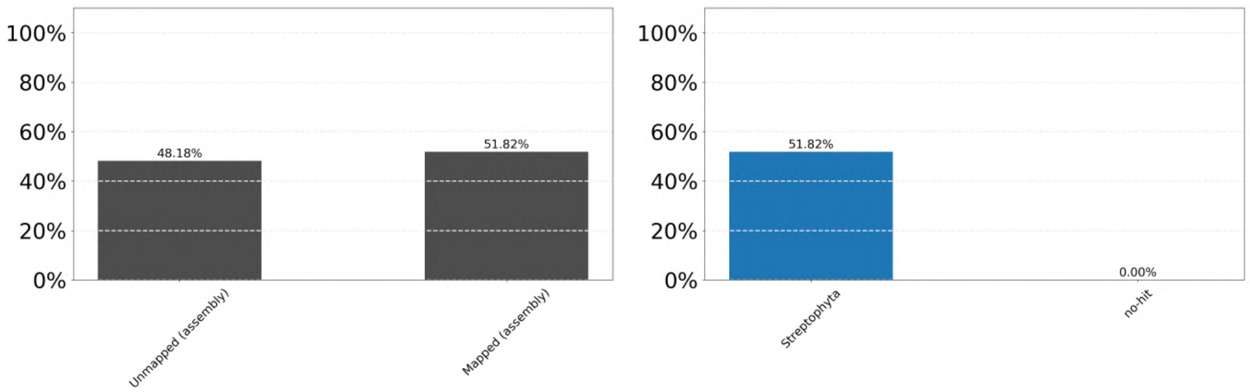

(B) Haplotype 1

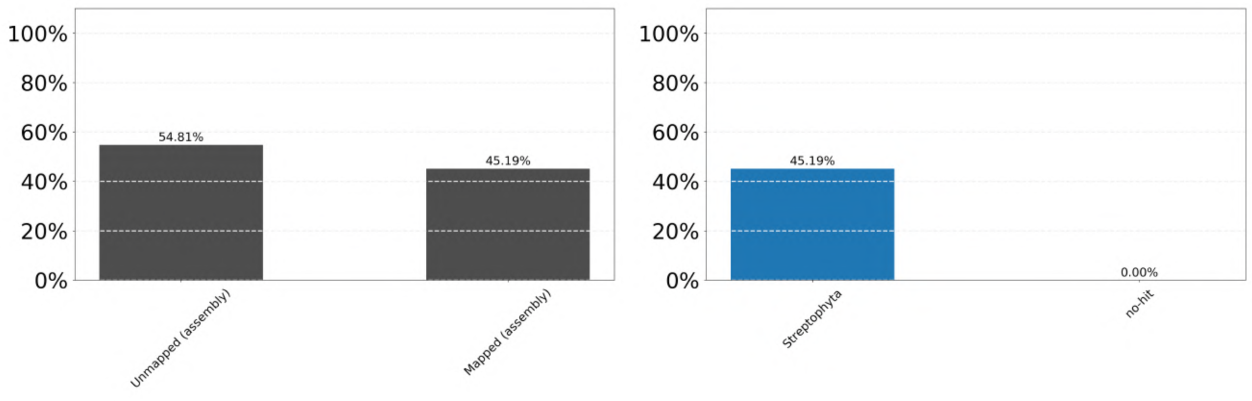


(C) Haplotype 2

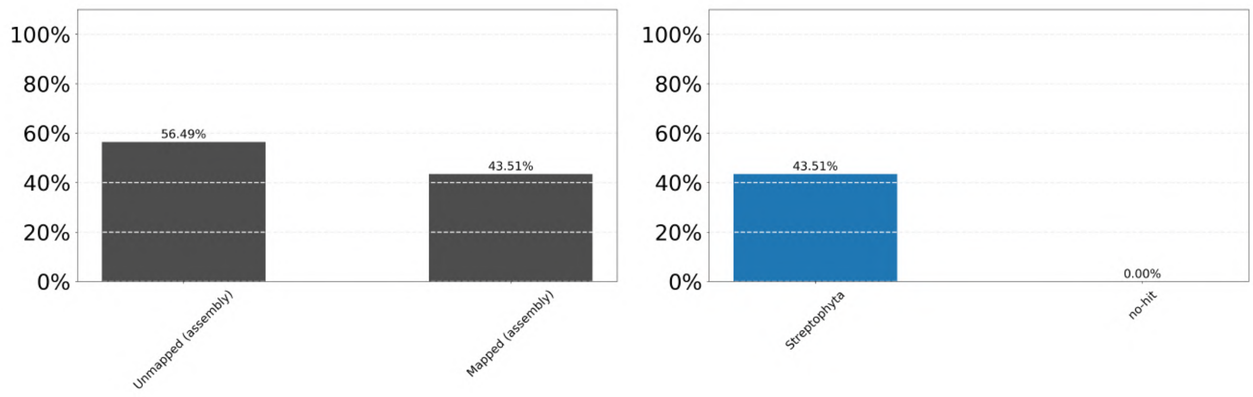


**Fig. S4:** Synteny visualization of the collinearity in the predicted proteins of the primary assembly.

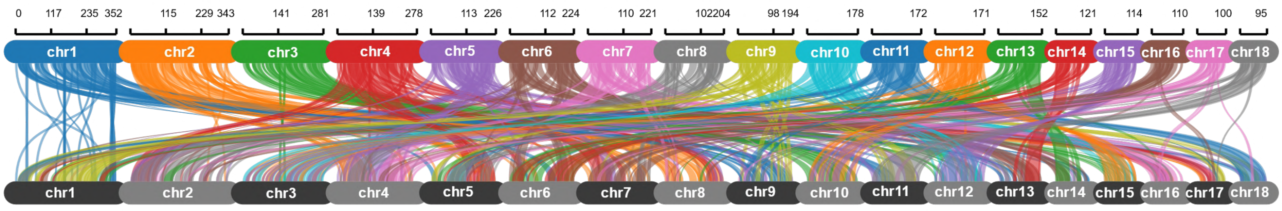

**Fig. S5:** Mitochondrial annotation obtained with MitoHifi for the primary genome assembly.

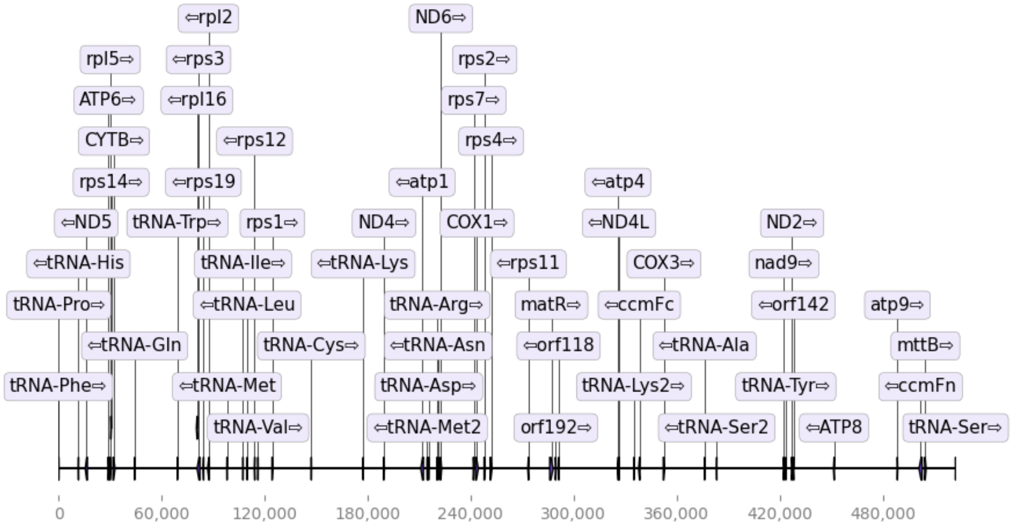

**Fig. S6:** Organelle genome annotation for (A) mitochondria and (B) chloroplast. The gray variation area in the inner circle indicates the GC content of the organelle genome.

(A) Mitochondrial genome

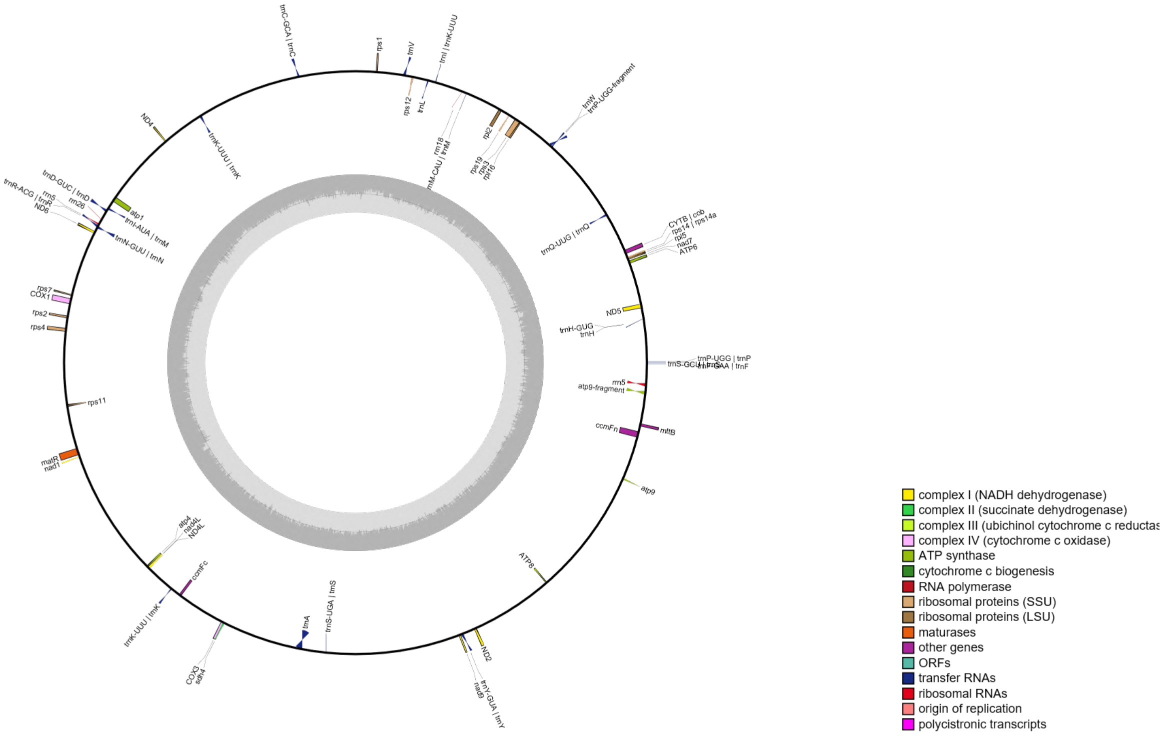


(B) Chloroplast genome

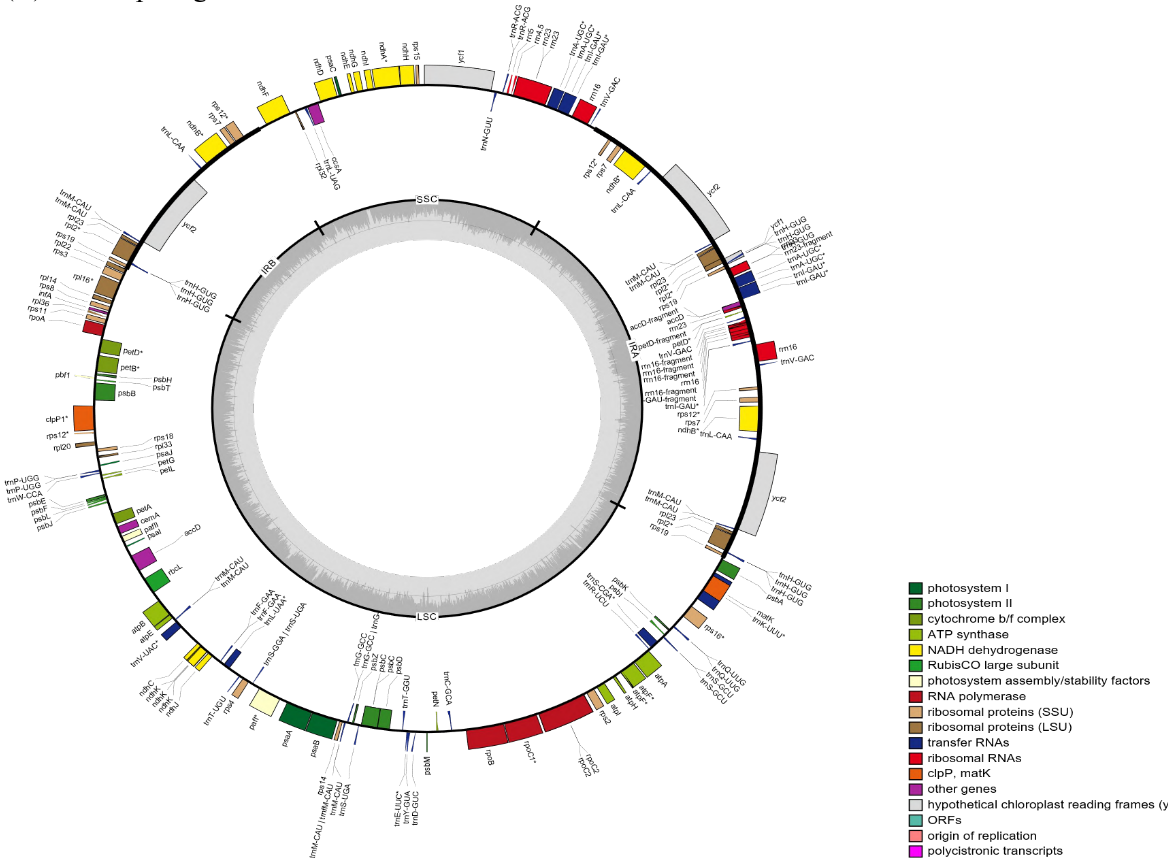

**Fig. S7:** Phylogenomic BUSCO tree of all published genomes. Light blue highlights complete and single-copy genes, dark blue represents complete and duplicated genes, yellow represents fragmented genes, and red represents missing genes.

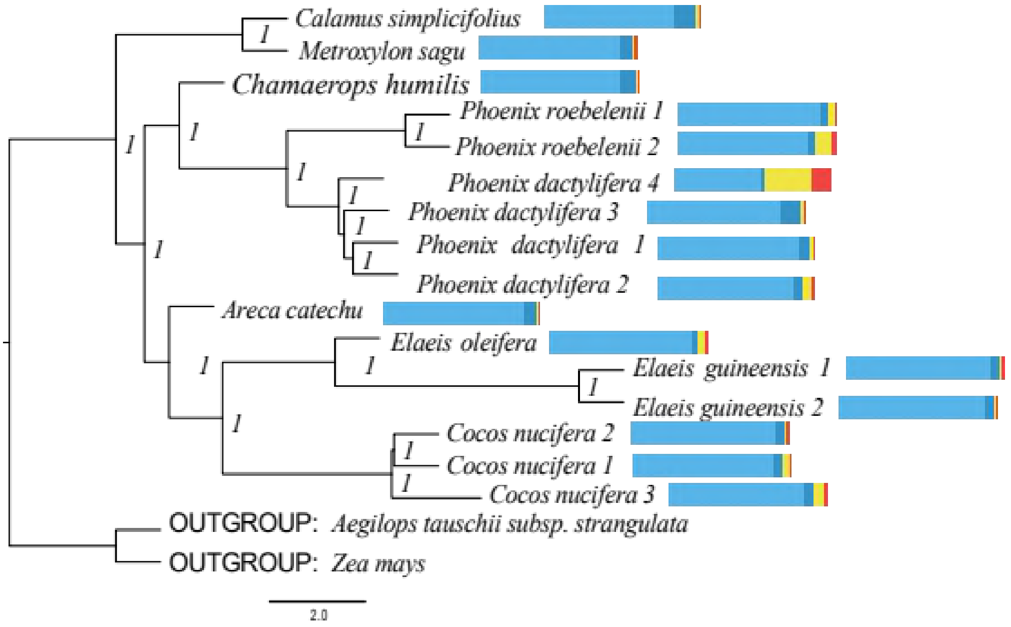


**Fig. S8:** Repeat divergence landscapes in several species of the family Arecaceae: (A) *Areca catechu,* (B) *Calamus simplicifolius,* (C) *Cocos nucifera,* (D) *Elaeis guineensis*, (E) *Elaeis oleifera*, (F) Metroxylon sagu, (G) *Phoenix dactylifera*, (H) *Phoenix roebelenii*, and (I) *Chamaerops humilis.*

(A) *Areca catechu*

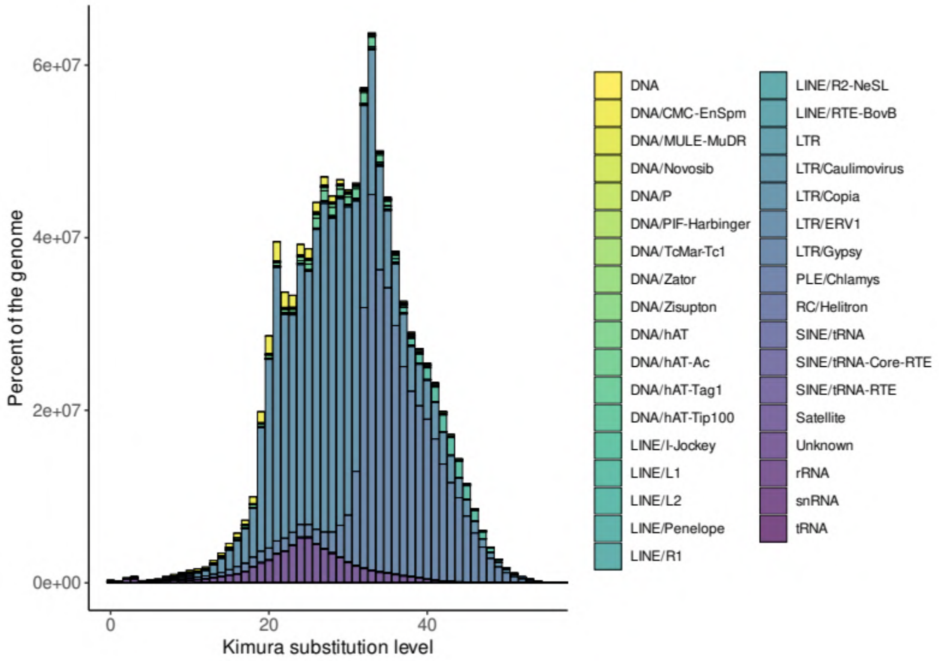

(B) *Calamus simplicifolius*

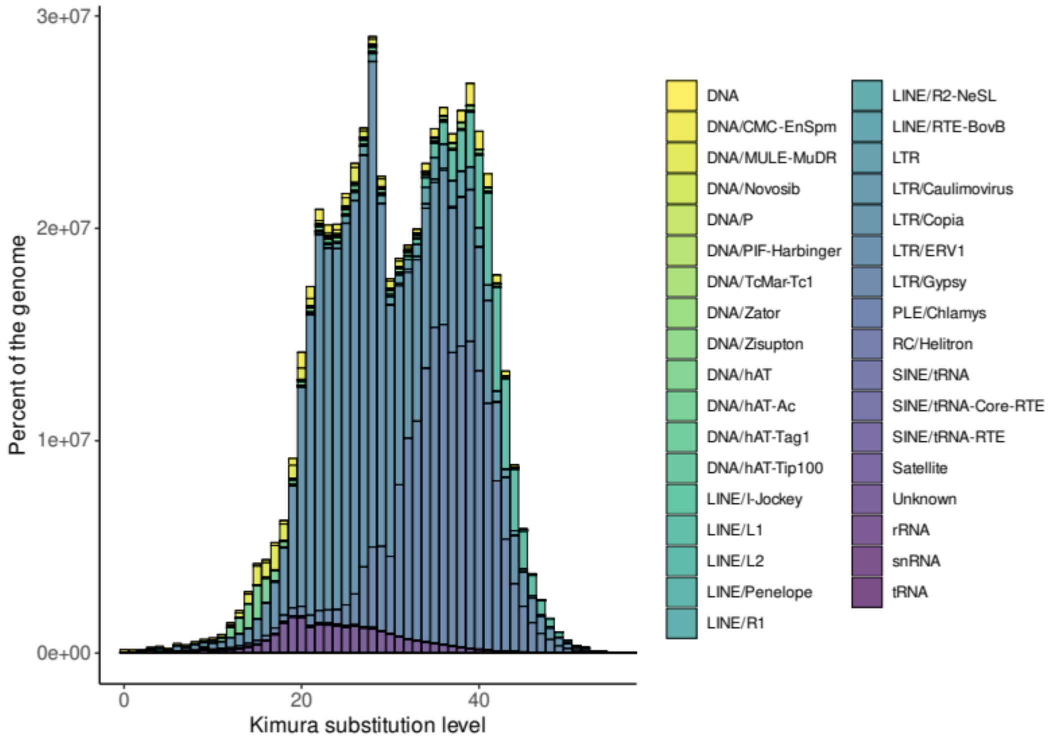


(C) *Cocos nucifera*

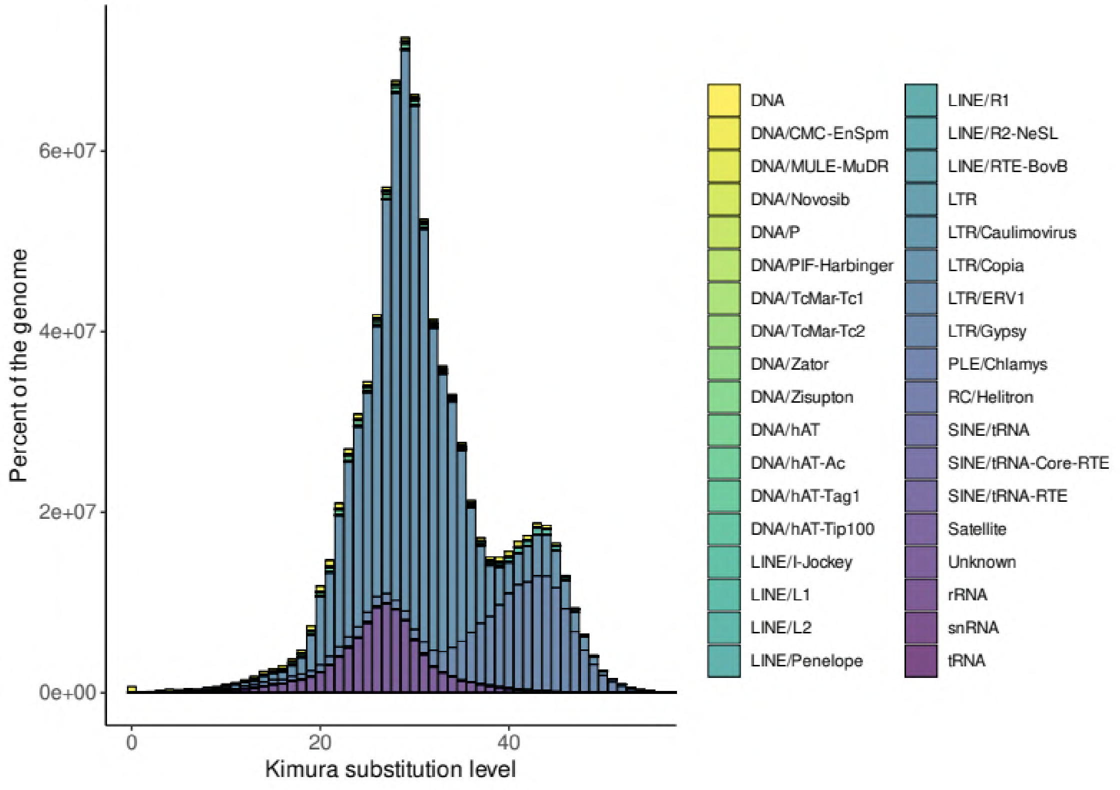

(D) *Elaeis guineensis*

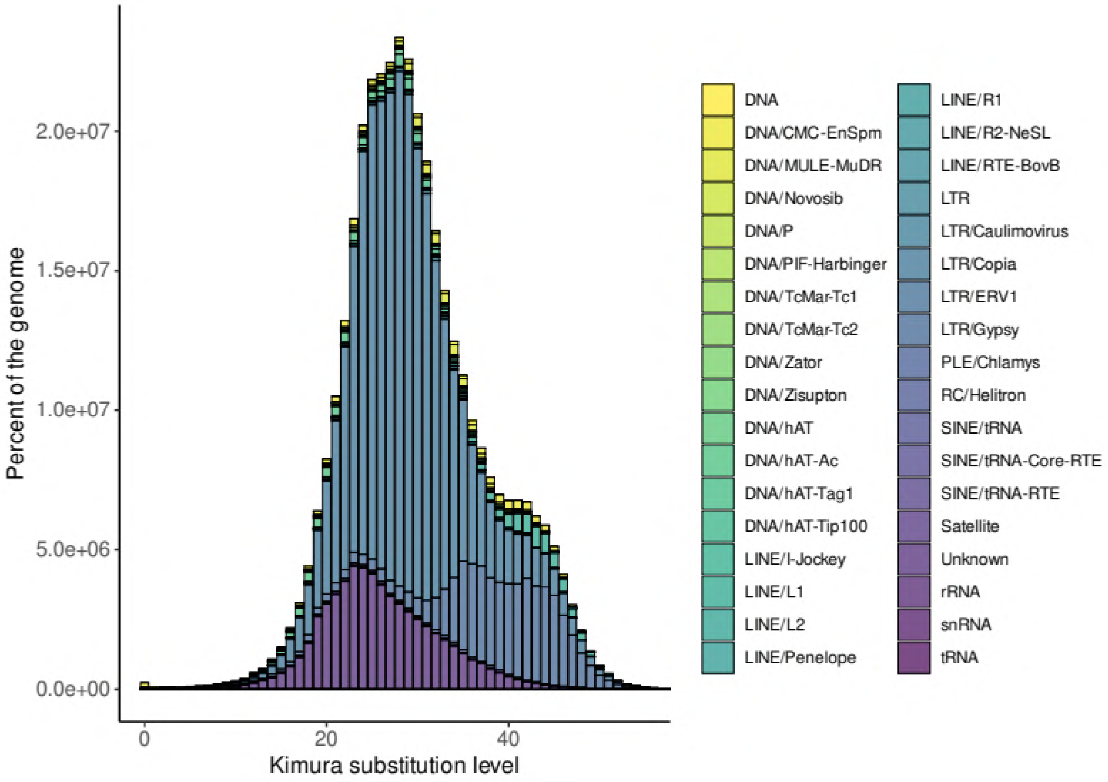


(E) *Elaeis oleifera*

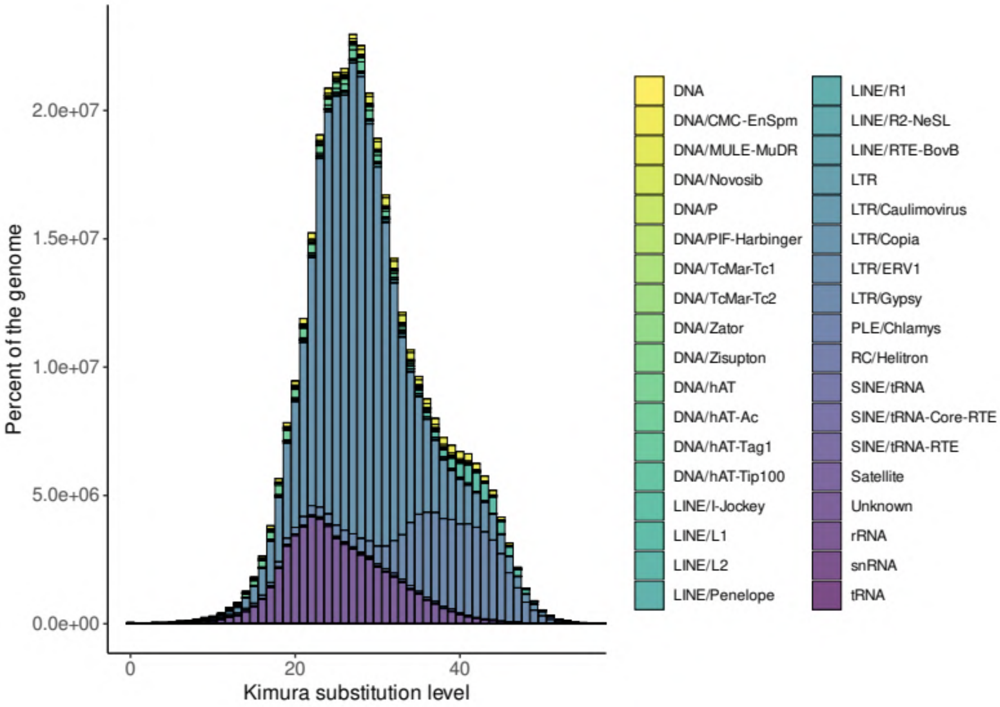

(F) *Metroxylon sagu*

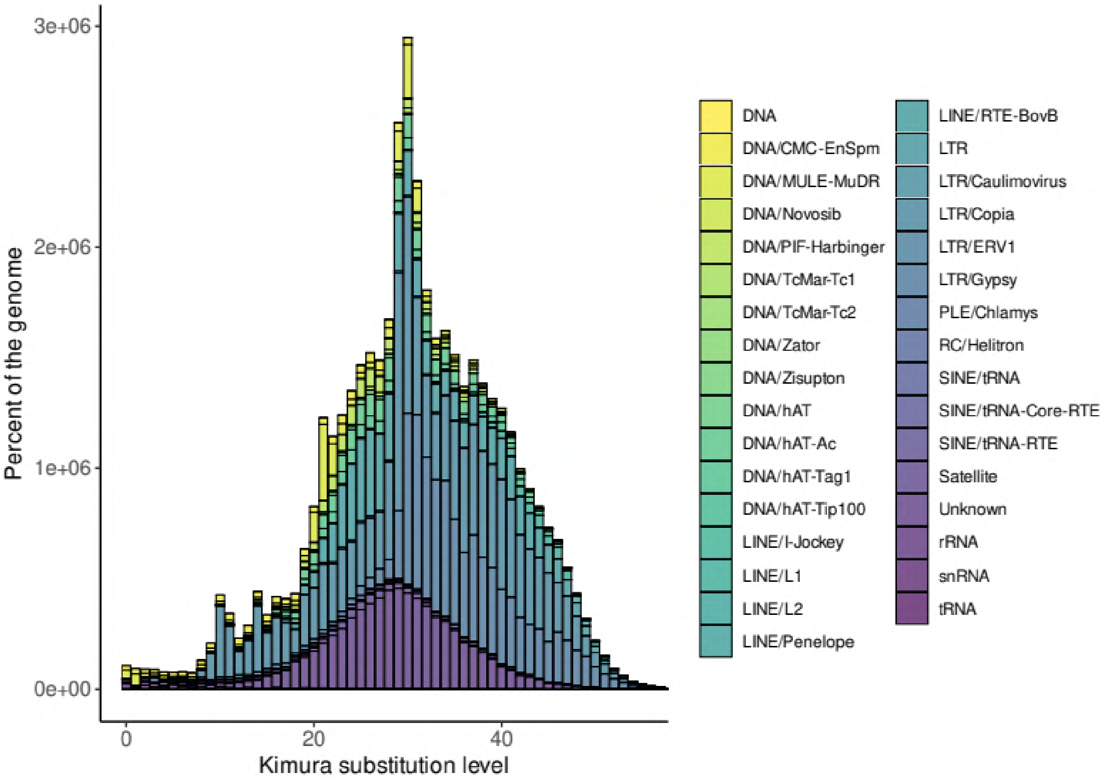


(G) *Phoenix dactylifera*

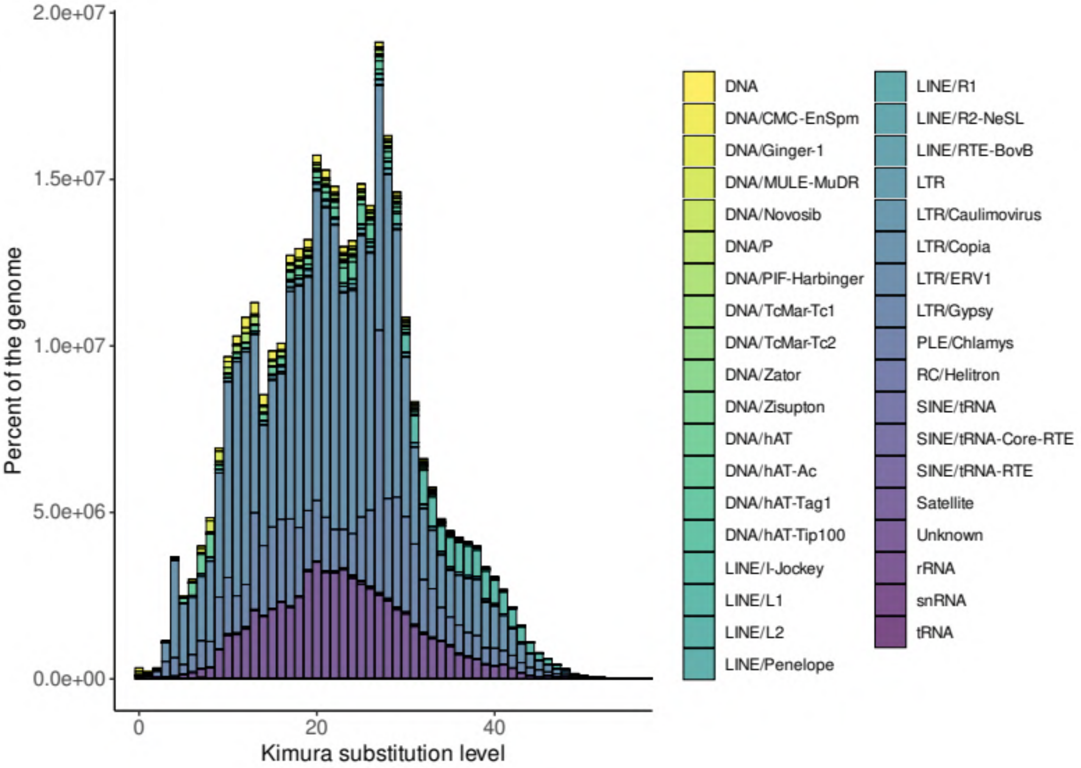

(H) *Phoenix roebelenii*

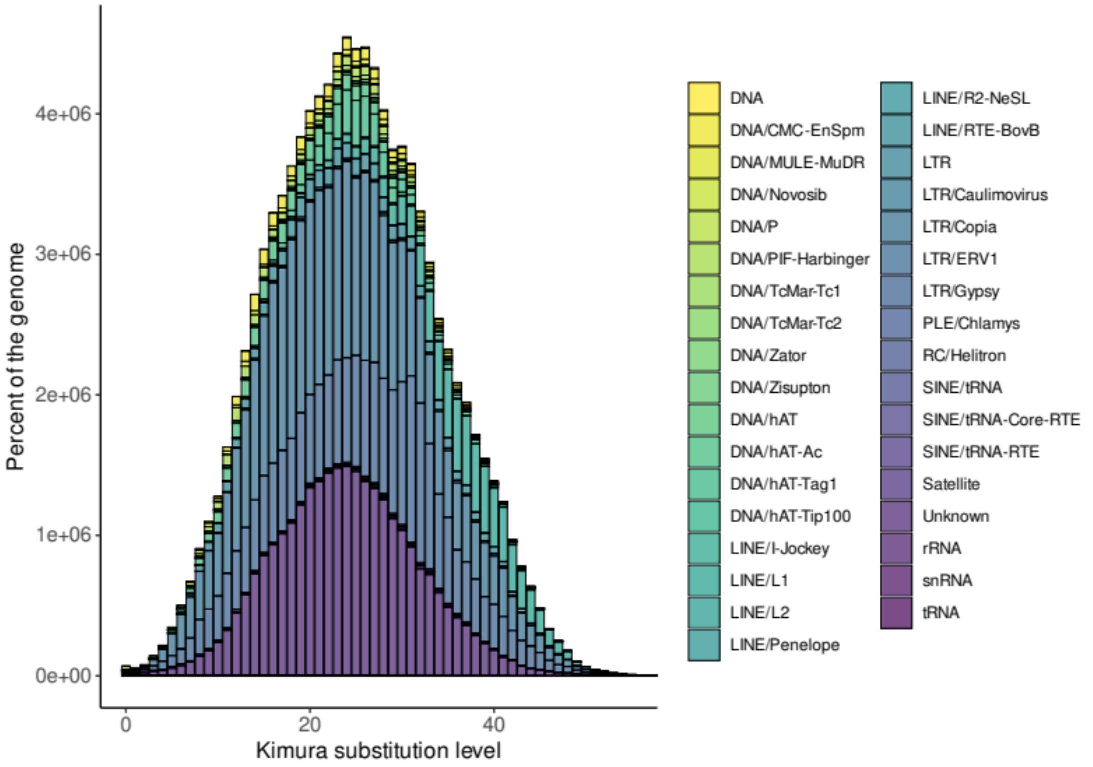


(I) *Chamaerops humilis*

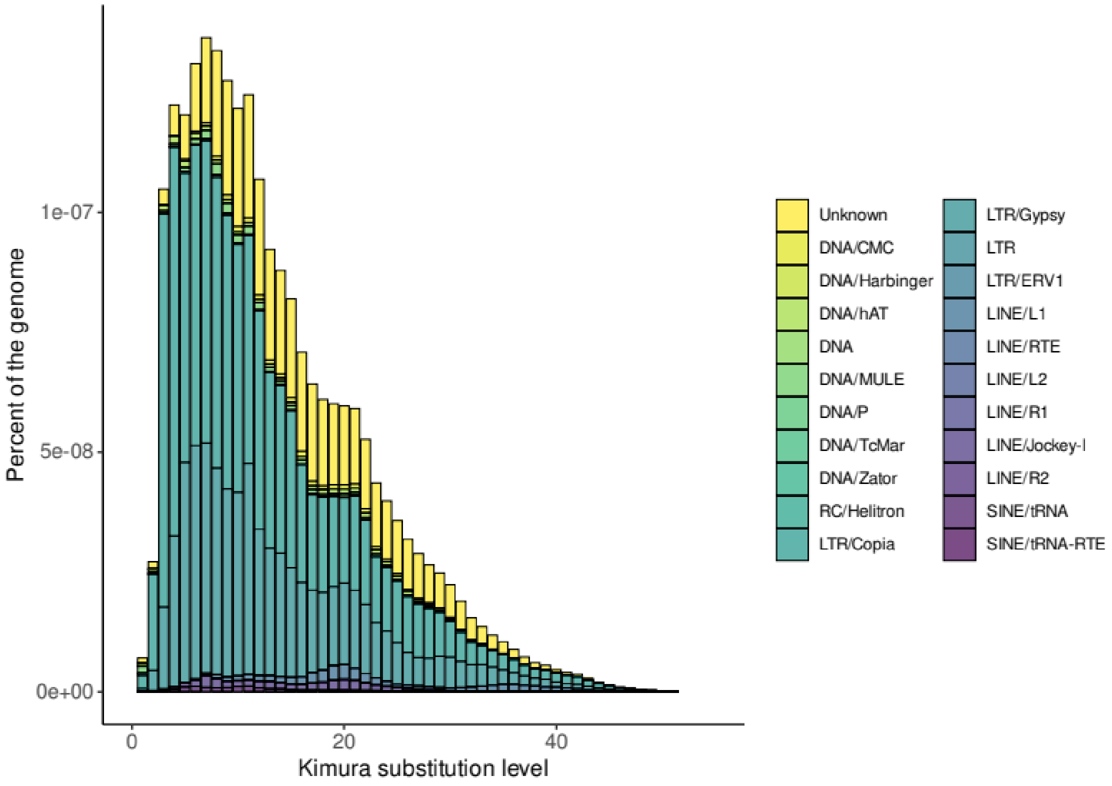
